# Modelling between-cell heterogeneity in within-host influenza virus infection

**DOI:** 10.64898/2026.05.17.725795

**Authors:** Ada W. C. Yan, Steven Riley, James M. McCaw

## Abstract

Cell tropism, or the preference of a virus for particular cell types, has major implications for viral transmission, pathogenesis, and evolution. An increase in viral fitness — increased within-host replication, also leading to increased transmission between hosts — can result from a virus changing its cell tropism. This is illustrated in the context of influenza, where adaptation to infect cells expressing ***α***2-6 linked sialic acid receptors enhances human-to-human transmissibility. Target cell populations differ not only in abundance but also in intrinsic properties such as susceptibility, viral production, and interferon responses, rendering the relationship between tropism and viral fitness multi-faceted and complex. Understanding how different cell tropisms quantitatively change fitness remains an important open question in virology and quantitative biology.

Here, we present a within-host mathematical model that incorporates distinct target cell types differing in key properties, and examine how cell tropism affects viral fitness, as measured by metrics such as peak viral load, infection duration, or total virus produced. Our analysis reveals that tradeoffs may arise when cell types differ by multiple characteristics. We further demonstrate that model parameters describing heterogeneity between cell types can be more accurately inferred when cell type proportions are measured alongside viral load. Our findings provide a framework for assessing the links between viral evolution, cell tropism, and within-host fitness, and motivate the design of experiments to collect quantitative data on between-cell heterogeneity.

## 1 Introduction

When a virus infects a host, different cell types are available to be infected. These cell types can differ in their susceptibility to infection, the amount of (viable and non-viable) virus they produce, production of interferon, and ability to enter an antiviral state. Cell tropism refers to the preference of a virus to infect one cell type over another. In some important cases, cell tropism is key to the fitness of a virus, defined as the ability of a virus to replicate within a host and cause onward transmission. For example, influenza viruses that cause pandemics have historically been of avian origin (with swine as an intermediate host in the case of the 2009 pandemic virus). In order to efficiently transmit between humans, influenza viruses need to evolve to preferentially infect cells which express *α*2-6 linked sialic acid receptors, rather than cells which express *α*2-3 linked sialic acid receptors that predominate in the reservoir species (Lipsitch et al. 2016). Non-ciliated cells, which express *α*2-6 linked sialic acid receptors, are more abundant in the human upper respiratory tract compared to ciliated cells which express *α*2-3 linked sialic acid receptors. For example, in the nasal cavity, only 20% of cells are ciliated, whereas in the bronchi branching into the lower respiratory tract, there are up to 90% ciliated cells (Yoshida et al. 2022; Sikkema et al. 2023). Therefore, it is plausible that a fitness advantage is conferred simply by adapting to infect more abundant cells at the initial site of infection. However, these cell types may also behave differently upon infection, such that the relationship between cell tropism and fitness becomes more complicated. For example, Roach et al. (2024) showed that when cell cultures with a mixture of cell types were infected with the same human influenza virus, a higher proportion of ciliated cells was associated with increased virus production. This result appears to contradict the observation that human influenza viruses must be adapted to infect non-ciliated cells, and suggests that the fitness advantage conferred by cell tropism could be more complicated, possibly due to cell-type-specific pro- or antiviral factors such as the innate immune response.

We wish to understand from a quantitative perspective: how do changes in viral fitness result from the evolution of viruses to infect different cell types? Given the characteristics of different cell types, is there always an optimal cell type to infect, or can there be tradeoffs such that infecting a similar proportion of each cell type is the optimal strategy? Many experiments in the scientific literature have been conducted where cells have been harvested from *in vitro* or *in vivo* infections, and analysed using techniques such as single-cell RNAseq (Fiege et al. 2021; Kelly et al. 2022; Lindeboom et al. 2024; Yoshida et al. 2022; Sikkema et al. 2023; Woodall et al. 2024) or flow cytometry (Roach et al. 2024). We would like to use these data to answer the proposed questions.

Within-host mathematical models have been used to quantify mechanisms of infection and immunity, and understand how these mechanisms contribute to the time course of infection. A number of mathematical models incorporating different types of target cells (cells available for infection) have been constructed and analysed in the literature. Many of these models are for chronic infection with pathogens such as HIV, so the focus of these analyses have been on the stability of equilibria (Elaiw 2010). Stability analysis for models of acute infection with multiple equilibria have also been studied (Payne et al. 1992; Wang et al. 2016). However, there are a small number of studies of the transient dynamics of infections (Perelson et al. 1997; Dahari et al. 2005; Bajaria et al. 2002). Some models include different spatial locations for infection (Higgins et al. 2025) and model the spread of virus between locations.

An important problem spanning many such models, and a ubiquitous challenge for within-host mathematical modelling, is that these models have many parameters, and it is unclear that their values can be estimated accurately from data.

In this study, we formulate a within-host viral dynamics model where cell types differ in their susceptibility to infection, amount of virus produced, and/or production of interferon. We show that changing the relative susceptibility to infection of the different cell types can lead to changes in fitness measured as measured by different metrics. We show that there can be tradeoffs in the optimal cell type to infect, for example if the cell type that produces the most virus also produces the most interferon. We then show that model parameters related to the degree of heterogeneity can be inferred if data on the proportion of cells of each type are available in addition to viral load data.

## 2 Methods

### 2.1 Model

We consider a simple model for acute viral infections, incorporating innate and adaptive immunity:

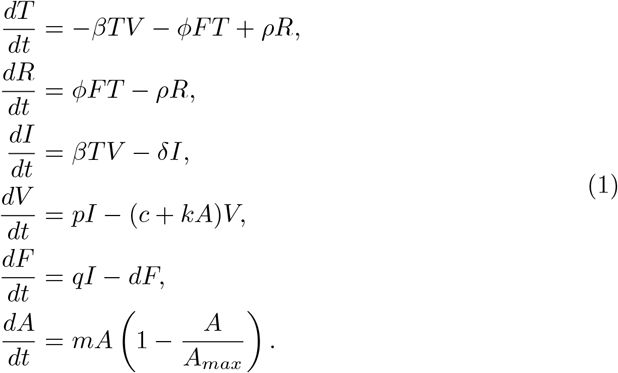

with initial conditions *T* (0) = *T*_0_, *R*(0) = 0, *I*(0) = 0, *V* (0) = *V*_0_, *F* (0) = 0, *A*(0) =*A*_0_.

This model draws upon models by Cao et al. (2015) and Handel et al. (2007). It includes the following processes:

- Target cells (*T*) become infected at rate *βV*;
- infected cells (*I*) produce free virus (*V*) at rate *p*;
- infected cells produce interferon (*F*) at rate *q*;
- free virus is neutralised by antibodies (*A*) at rate *k*;
- interferon renders target cells resistant to infection (*R*) at rate *ϕ*;
- resistant cells revert to being susceptible at rate *ρ*;
- virus, infected cells, and interferon decay at rates *c, δ* and *d* respectively;
- antibodies are not modelled mechanistically, but they are assumed to grow exponentially at rate *m* from the start of infection onwards, with a carrying capacity *A*_*max*_.

Note that *V* refers to free infectious virus (e.g. measured by TCID_50_ or plaque assay) rather than genome copy number (e.g. measured by RT-qPCR). While many different mathematical forms for the innate and adaptive immune responses in influenza virus infection have been proposed in the literature (e.g.Baccam et al. (2006); Smith et al. (2018); Pawelek et al. (2012); Cao et al. (2015); Miao et al. (2010)), we chose simple forms that are sufficient for the purposes of our study. Some immune mechanisms such as natural killer cells and CD8+ T cells have not been included. The model structure and chosen parameter values are such that the innate immune response prevents target cell depletion and lowers the peak viral load, and that the adaptive immune response, rather than target cell depletion, is responsible for resolution of the infection.

With *N* cell types, the model equations become

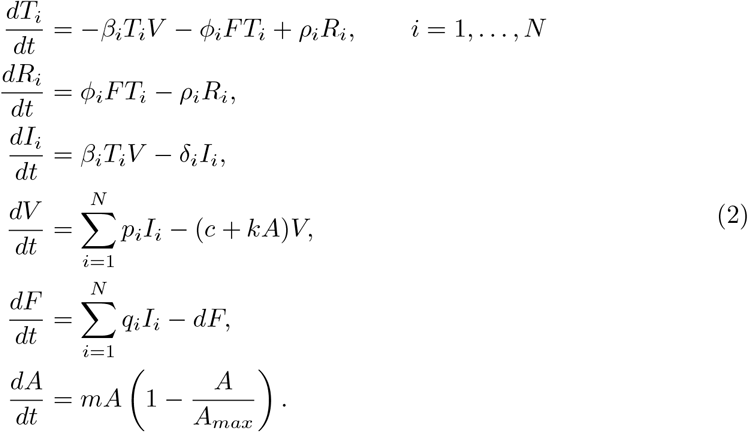

with initial conditions *T*_*i*_(0) = *T*_0*i*_, *R*_*i*_(0) = 0, *I*_*i*_(0) = 0, *V* (0) = *V*_0_, *F* (0) = 0, *A*(0) = *A*_0_. Figure 1 shows the compartmental model in Eq. 2 for *N* = 2 cell types.

**Fig. 1:**
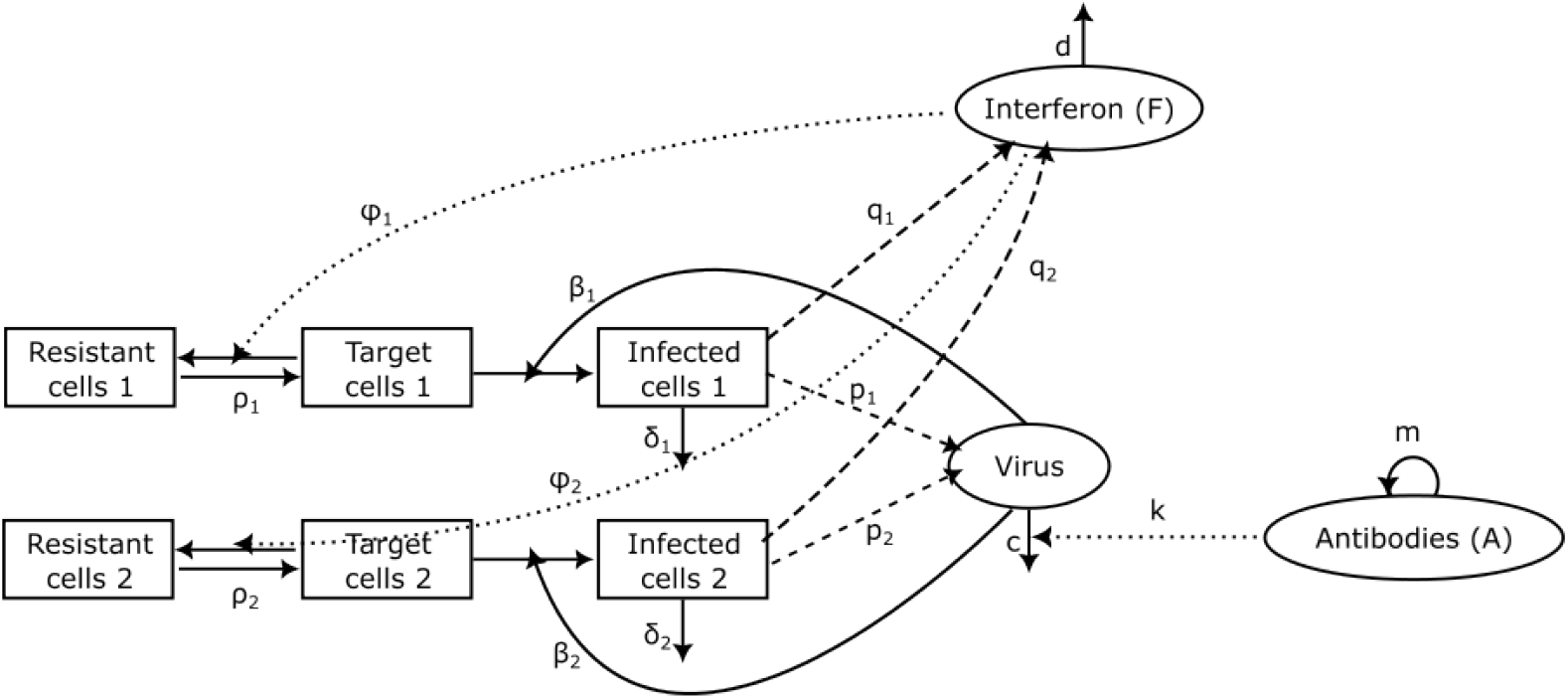
Compartmental model in Eq. 2 for *N* = 2 cell types.

Table 1 shows the baseline values for each parameter, based on those by Cao et al.(2015) and Handel et al. (2007). The parameter values were chosen such that the resulting viral load trajectories resembled those observed for *in vivo* influenza virus infections in ferrets. Specifically, the parameter values for *β, δ, p, ϕ, ρ, q, T*_0_ and *V*_0_ were taken directly from Cao et al. (2015). The antibody parameters *m, A*_0_ and *A*_*max*_ and viral decay parameter *c* were tuned such that the simpler mathematical form of the adaptive immune response produced similar quantitative results to Cao et al. (2015). Lack of parameter identifiability for viral dynamics models has previously been documented (Yan et al. 2019); there could potentially be other choices of parameter values producing realistic viral load curves, but we have chosen a single plausible parameter set for illustration of the model behaviour. Interferon and antibodies were quantified in relative units *u*_*F*_ and *u*_*A*_ as these compartments are often not directly measured over time.

**Table 1.** Baseline parameter values.

| Parameter | Value | Unit |
| --- | --- | --- |
| $\beta$ | $5 \times 10^{-5}$ | $\text{TCID}_{50}^{-1} \text{d}^{-1}$ |
| $\delta$ | 3 | $\text{d}^{-1}$ |
| $p$ | 0.35 | $\text{TCID}_{50} \text{cell}^{-1} \text{d}^{-1}$ |
| $c$ | 4 | $\text{d}^{-1}$ |
| $\phi$ | 0.14 | $u_F^{-1} \text{d}^{-1}$ |
| $\rho$ | 0.05 | $\text{d}^{-1}$ |
| $q$ | $5 \times 10^{-5}$ | $u_F \text{cell}^{-1} \text{d}^{-1}$ |
| $k$ | 1 | $u_A^{-1} \text{d}^{-1}$ |
| $m$ | 1 | $\text{d}^{-1}$ |
| $T_0$ | $7 \times 10^7$ | cell |
| $V_0$ | 1 | $\text{TCID}_{50}$ |
| $A_0$ | 0.34 | $u_A$ |
| $A_{max}$ | $10^4$ | $u_A$ |

### 2.2 Viral load simulation and fitness metrics

We used `odin` 1.2.6 (FitzJohn 2024) in `R` 4.4.0 (R Core Team 2024) to numerically solve Eq. 2 for *N* = 2 cell types, changing the amount of heterogeneity in one or more parameters. The model captures cell tropism through having different susceptibilities to infection (*β*_*i*_) for each cell type, so *β*_*i*_ was varied for all simulations.

We also constructed metrics to summarise viral fitness into quantities more readily comparable between viruses: the within-host basic reproduction number (*R*_0_); the growth rate (*r*); and a third metric designed to capture the overall level of transmission to another host. Both the within-host basic reproduction number (*R*_0_) and growth rate (*r*) affect the ability of a virus to compete with a co-infecting virus within the same host. The basic reproduction number and growth rate can be calculated using the next generation matrix approach (Diekmann et al. 2010), linearising the ODEs around the disease-free equilibrium:

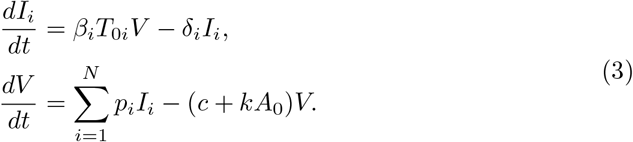

The basic reproduction number for a within-host viral dynamics model is conventionally defined as the mean total number of virions produced by secondary infected cells resulting from a single initial virion in an pool of initially uninfected cells:

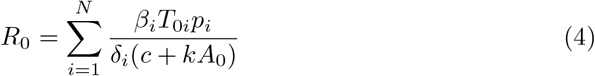

We note that the next generation matrix approach defines the time between introducing the first virion, through to infection of a cell, and finally for a secondary virion to be produced, to be two generations. Thus, the *R*_0_ calculated using the next generation matrix needs to be squared to yield the result in Eq. 4.

In the case that all *δ*_*i*_ = *δ*, i.e. the infected cell death rate is the same across all cell types, the growth rate is

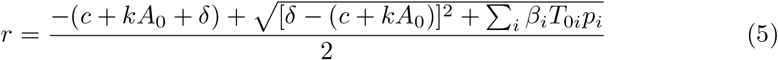

The third metric, related to the amount of virus shed by a host, affects the ability of a virus to transmit to another host. A number of different mathematical forms for this metric have been adopted in the literature, as compared by Asher et al. (2023). For our primary analysis we consider the area under the viral curve (AUC), defined as

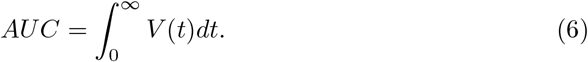

We also consider a number of alternative relationships to map from viral kinetics to transmission (described and presented in the sensitivity analysis in Sec. 3.3).

### 2.3 Simulation estimation study

We conducted a simulation-estimation study to investigate whether the degree of target cell heterogeneity can be inferred given different types of data, and how the identifiability of parameters with heterogeneity impacts the ability to make accurate model predictions. The number of combinations of parameters that could vary between cell types is large; we restricted the scope of this study to the case where the cell types differ in susceptibility (*β*_*i*_) and abundance (*T*_0*i*_).

#### 2.3.1 Generating parameter sets for simulation

For a model with two cell types (*N* = 2) and heterogeneity in susceptibility to infection (*β*_*i*_) only, to improve exploration of parameter space, we reparameterised *β*_*i*_ in terms of the weighted mean of *β*_*i*_, which we denote 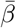, and a parameter Δ which ranges between [− 1, 1] and controls the degree of heterogeneity between the two cell types. The equation for 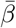 is:

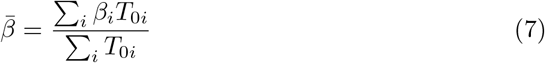

For the homogeneous model with *N* = 1, we have shown in previous work that the initial growth rate *r* is identifiable (Yan et al. 2019). Eq. 5 shows that when there is only heterogeneity in *β*_*i*_ and *T*_0*i*_, *r* is a function of Σ_*i*_ *β*_*i*_*T*_0*i*_, rather than *β*_*i*_ and *T*_0*i*_ independently. Thus we expect 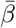 to be identifiable in a simulation-estimation study, justifying our choice of re-parameterisation.

We implicitly define Δ by the following functions to calculate *β*_*i*_ given 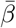 and Δ:

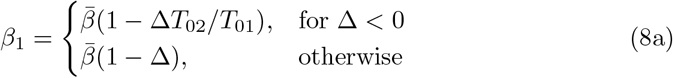

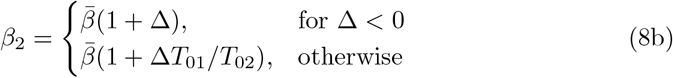

This definition ensures that *β*_1_ = *β*_2_ for Δ = 0; *β*_2_ = 0, 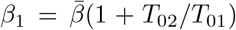 for Δ = −1; and 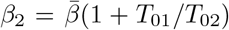 for Δ = 1. When Δ is thus defined, then when its value is 0, setting *β*_1_ = *β*_2_ and solving the ODEs in Eq. 2 leads to the same viral load as the homogeneous model, regardless of the values of *T*_0*i*_ (assuming all other parameters are the same between compartments). The natural bounds for this definition of Δ (−1 and 1 when *β*_1_ or *β*_2_ are 0 respectively) are also the same for all values of *β*_1_ and *β*_2_. This definition thus enables us to compare the effects of heterogeneity across different parameter sets.

First, we used Latin hypercube sampling (implemented in *lhs* 1.2.0 (Carnell 2024)) to generate 5000 parameter sets, varying 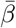, *δ, p/δ, c* and *V*_0_ in log space by one order of magnitude above and below the values given in Table 1, and *T*_01_*/T*_0_ in linear space between 0 and 1. The remaining parameters were held fixed. From the 5000 parameter sets, we selected those with realistic initial growth rates *r* (based on the homogeneous case, see Eq. 5) in the range ln(10^3^) ≤*r* ≤ln(10^5^). As *V* (*t*) ≈ *V* (0) exp(*rt*) in the initial phase of the infection, this restriction ensures that the viral load increases by 3–5 orders of magnitude during the first day of infection. 100 parameter sets (out of 558) fulfilling these criteria were randomly selected for subsequent analysis.

#### 2.3.2 Simulating data

For each of the 100 parameter sets, we solved the model system equations (Eq. 2) and generated simulated data for Δ = −0.8, Δ = 0, and Δ = 0.8. Simulated viral load data and cell proportion data were generated in triplicate for each parameter set.

We assume that cell proportion data are generated using common techniques such as flow cytometry or single-cell RNAseq. With these techniques, it is possible to distinguish infected from uninfected cells (using virus protein-specific antibodies for flow cytometry or proportion of viral RNA in a cell for single-cell RNAseq), and to distinguish different cell types (using antibodies for cell type markers for flow cytometry or cell type annotation for single-cell RNAseq). However, distinguishing between uninfected cells which are susceptible to infection (*T*) versus resistant to infection (*R*) is difficult. Hence, we define a variable for the number of uninfected cells, *U*_*i*_(*t*) = *T*_*i*_(*t*) + *R*_*i*_(*t*). Then, we generate cell proportion data by calculating the proportion of uninfected and infected cells of each type at a given time, out of total live cells, i.e., the data at time *t*_*i*_ are

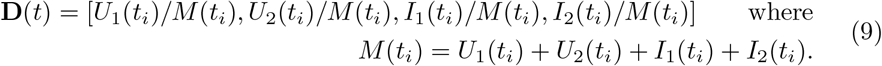

Cell proportion data were sampled on days 1 and 2 as the experimental methods to sample these data destroy the sample, limiting the number that can be taken. Each sample consists of a subset of the live cells at that time. We assume a sample size of 5000. The cell proportion data are assumed to follow a Dirichlet multinomial distribution with overdispersion parameter *ρ*_0_ = 0.01, that is [*Û*_1_(*t*_*i*_), *Û*_2_(*t*_*i*_), *Î*_1_(*t*_*i*_), *Î*_2_(*t*_*i*_)] ~ DirMult(*C, α*) where *α* = [*U*_1_(*t*_*i*_)*/M, U*_2_(*t*_*i*_)*/M, I*_1_(*t*_*i*_)*/M, I*_2_(*t*_*i*_)*/M*] × (1 − *ρ*_0_)*/ρ*_0_.

Viral load was sampled daily for seven days, with lognormal noise added to each replicate, such that 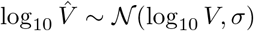 where *σ* = 0.5. An observation threshold was then imposed such that values under the lower limit of detection (10 TCID_50_) were censored.

#### 2.3.3 Estimating parameters

Once we sampled “true” parameter sets and simulated data using these parameter sets, we re-estimated the parmaeters from the simulated data to see if we could recover the true values. Inference was conducted using the No U-Turn sampler in `Stan` ({Stan Development Team} 2025) 2.37.0, with the interface `cmdstanr` (Gabry et al. 2025) 0.8.1. Using the default settings, eight parallel chains were sampled, with 1000 warm-up iterations and 1000 sampling iterations each. We expected some parameters to be unidentifiable, which is known to cause issues with convergence of chains, as well as divergent transitions in the No U-Turn sampler. We removed chains where divergent transitions were observed and chose a subset of chains (at least three), such that we retained as many chains as possible while keeping 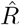, a measure used to assess convergence, below 1.01. If this was not possible, we dropped the requirement for chains to have no divergent transitions, and chose the subset of chains (at least three) which minimized 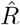.

We estimated *β* and Δ either using viral load data only, or using both viral load and cell proportion data. All other parameters were assumed to be known, as we are focusing on the identifiability of parameters modelled to have heterogeneity. We set the prior distribution for *β* as uniform in log space (log_10_ *β* ~ 𝒰 (−8, −4)), and the prior distribution for Δ as uniform between 0 and 1.

Code to reproduce all results in the study, and all simulated data, can be found at https://github.com/ada-w-yan/fluheterogeneitypaper.

## 3 Results

When a virus’ cell tropism changes and it results in increased fitness, this could be attributed to the higher abundance of the newly preferred cell type (Lipsitch et al. 2016); an increased viral production rate (for example, through more efficient use of host proteins in replication and/or transcription (Long et al. 2016)); or a decreased rate of innate immune response induction.

In Section 3.1, we first confirm whether the model reproduces expected changes in fitness under various scenarios. In Section 3.2, we then use the model to predict changes to viral fitness if more than one of these factors is at play. In Section 3.3, we investigate whether the changes observed in the transmissibility metric are sensitive to the mathematical form of the metric. Finally, we conduct a simulation-estimation study to determine the types of data required to estimate the degree of heterogeneity between cells.

### 3.1 The model captures increased fitness when preferentially infecting cells which are more abundant, produce more virus, or produce less interferon

We first confirm whether the model produces hypothesised behaviours of increased fitness for three scenarios:

A. preferentially infecting cells which are more abundant;
B. preferentially infecting cells which produce more virus;
C. preferentially infecting cells which produce less interferon.

In all scenarios in this study, we simulate a changing cell tropism by varying *β*_1_ and *β*_2_ (susceptibility to infection of each cell type) while keeping their sum, *β*_1_ + *β*_2_, fixed. That is, we assume that when a virus evolves to change the cell type it prefers to infect, it increases the rate at which it infects cell type 1 by the same amount it decreases the rate at which it infects cell type 2 (or vice versa). In reality, virus evolution could result in arbitrary changes to *β*_1_ and *β*_2_, rather than necessarily evolving along a trajectory where *β*_1_ + *β*_2_ stays constant. However, if we consider the case where cell types are identical and equally abundant (same *p*_*i*_, *δ*_*i*_, *q*_*i*_, *ϕ*_*i*_, *ρ*_*i*_ and *T*_0*i*_), then changing *β*_1_ and *β*_2_ while keeping *β*_1_ + *β*_2_ constant conserves *R*_0_ (by examination of Eq. 4). Thus, if *β*_1_ and *β*_2_ are changed while keeping their sum constant, any change in *R*_0_ must be due to heterogeneities between cell types. Conversely, if *β*_1_ and *β*_2_ are allowed to change arbitrarily, changes in *R*_0_ are partially attributable to a change in the mean susceptibility of cells had the cell types been otherwise identical and equally abundant. Thus, we focus on the scenario where *β*_1_ and *β*_2_ are varied while keeping their sum fixed.

In scenario A (unequal cell abundances), we set *T*_01_ = 5.6 × 10^7^, *T*_02_ = 1.4 × 10^7^, such that cell type 1 has an 80% abundance, similar to non-ciliated cells in the human upper respiratory tract (Yoshida et al. 2022; Sikkema et al. 2023) All other parameters are equal between cell types as per Table 1. For scenario B (unequal virus production rates), we set *p*_1_ = 0.56, *p*_2_ = 0.14, such that cell type 1 produces virus at a greater rate. Inferring production rates of virus from different cell types from existing data is less straightforward, so we have set that cell type 1 produces 80% of virus to keep the ratios the same as scenario A. All other parameters are equal between cell types. For scenario C (unequal interferon production rates), we set *q*_1_ = 2.5 × 10^−5^, *q*_2_ = 7.5 × 10^−5^, such that cell type 1 produces interferon at a lower rate, and all other parameters are equal between cell types. While some choices in the degree of heterogeneity were arbitrary, the results are intended to show possible qualitative behaviours of the model and are not tied to particular choices of parameter values.

Figure 2 shows the simulated viral loads for Scenarios A, B and C as *β*_1_ − *β*_2_ is changed (colours) while *β*_1_ + *β*_2_ is held constant. That is, we see how the change in cell tropism leads to a change in viral load if cells have unequal abundance, virus production rates are unequal between cells, or interferon production rates are unequal between cells. In Scenarios A and B, the viral load increases more quickly initially for viruses preferentially infecting cell types that are more abundant (Scenario A) or that produced more virus (Scenario B). The initial rate of increase of viral load is the same regardless of cell tropism in Scenario C where the cell types differed by induction rate of interferon. In all three scenarios, as expected, the peak viral load is higher for viruses preferentially infecting cell types that are more abundant (Scenario A), that produce more virus (Scenario B), or that produce less interferon (Scenario C). However, it is only in scenarios A and B that the initial growth rate is impacted by cell tropism. We now explore how this result relates to the change in within-host reproduction number.

**Fig. 2:**
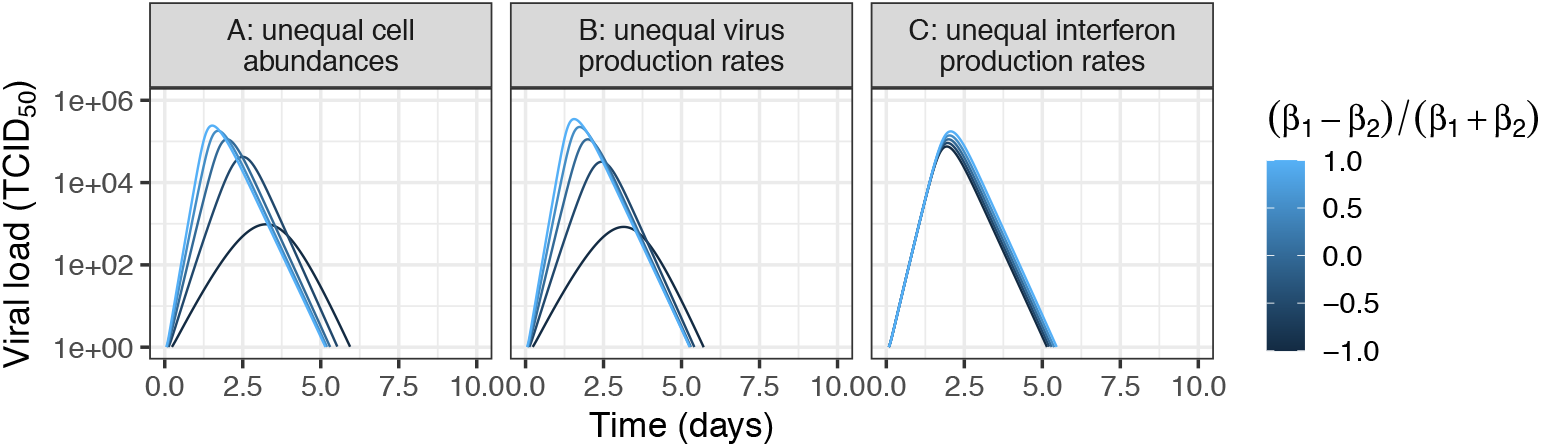
Simulated viral loads for scenarios A (unequal cell abundances), B (unequal virus production rates) and C (unequal interferon production rates) when *β*_1_ − *β*_2_ is changed (colours) while *β*_1_ + *β*_2_ is held constant.

Figure 3 shows (top) the within-host basic reproduction number and (bottom) the AUC for scenarios A, B and C when *β*_1_ − *β*_2_ is changed (x-axis) while *β*_1_ + *β*_2_ is held constant. For Scenarios A (preferentially infecting cell types that are more abundant) and B (preferentially infecting cell types that produce more virus), increasing preference for cell type 1 increases the basic reproduction number, while for Scenario C (preferentially infecting cell types that produce less interferon), the basic reproduction number remains constant regardless of cell tropism. As the definition of the basic reproduction number pertains to secondary virions starting from a completely susceptible population of cells with no interferon, the changes in eventual interferon levels due to changed cell tropism do not change the basic reproduction number.

**Fig. 3:**
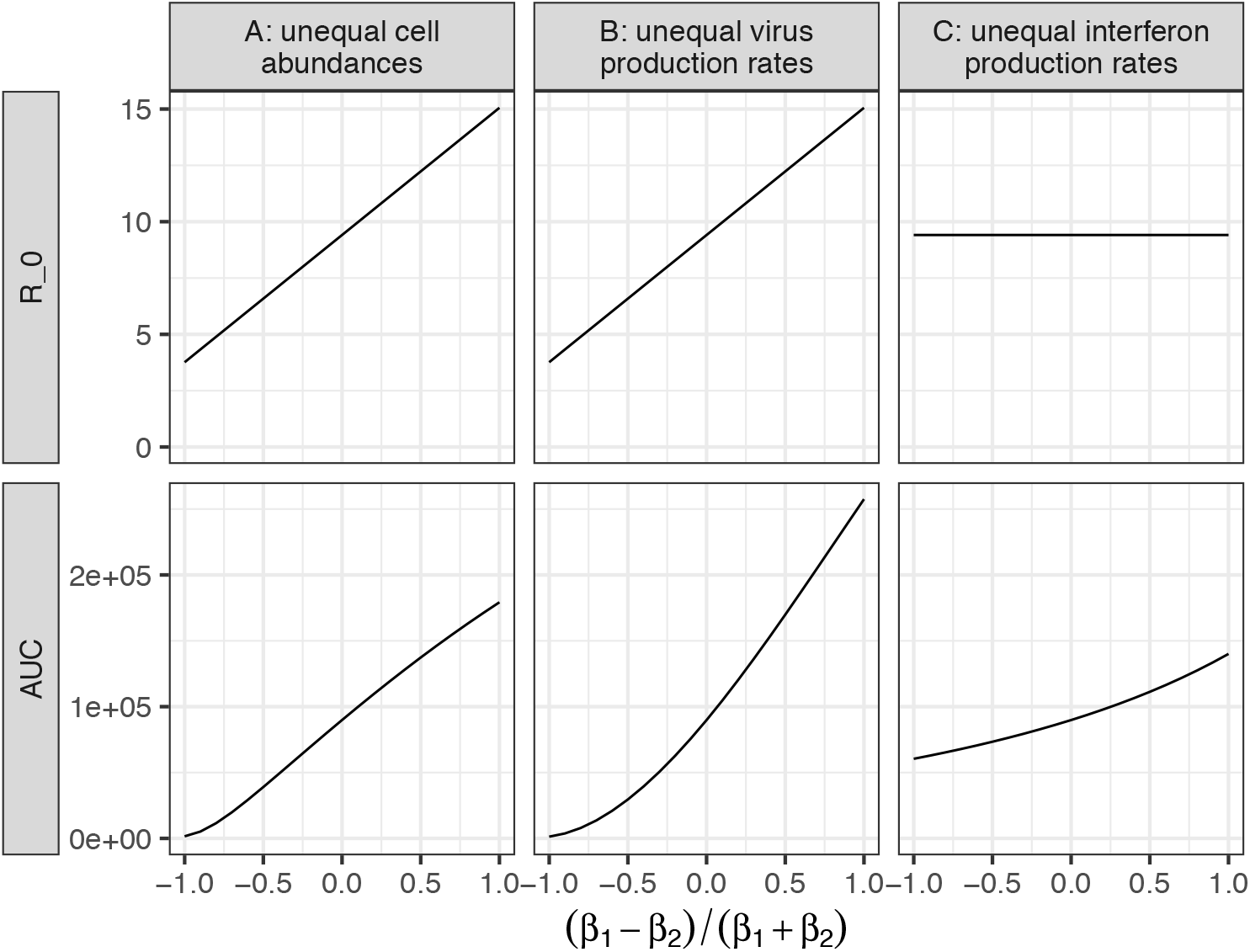
(Top) Basic reproduction number (bottom) AUC for scenarios A, B and C when *β*_1_ − *β*_2_ is changed (x-axis) while *β*_1_ + *β*_2_ is held constant.

For a model with equal *δ*_*i*_, we can multiply both sides of Eq. 4 by *δ*(*c* + *kA*_0_) and substitute into Eq. 5 to yield

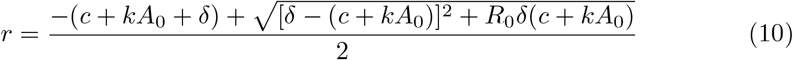

Thus, for fixed *c* + *kA*_0_ and *δ, r* changes monotonically with *R*_0_. In other words, a change in *β*_*i*_, *p*_*i*_ or *T*_0*i*_ which increases *R*_0_ also increases *r*, and vice versa. Therefore, the relationship between *β*_1_ − *β*_2_ and *R*_0_ also holds for *β*_1_ − *β*_2_ and *r*.

For all scenarios, an increasing preference for cell type 1 increases the area under the viral load curve.

### 3.2 Tradeoffs when cells differ in more than one aspect

So far, we have considered cases where cells differ in abundance, viral production rate, or interferon production rate, in addition to susceptibility to infections. A natural question to consider, both from first principles and as a logical consequence of the above two results, is what would happen if the cells differ in more than one aspect other than tropism?

First, we use the model to predict how viral fitness changes with cell tropism when the cell type with a higher viral production rate is also (relatively) less abundant than the other cell type (Fig. 4A). We fixed *T*_01_ = 2.1 *×* 10^7^, *T*_02_ = 4.9 *×* 10^7^ and simulate using three different sets of values for *p*_*i*_: i) *p*_1_ = 0.42, *p*_2_ = 0.28; ii) *p*_1_ = 0.49, *p*_2_ = 0.21; iii) *p*_1_ = 0.63, *p*_2_ = 0.07 TCID_50_cell^−1^d^−1^. For all three of these parameter sets, *p*_1_ > *p*_2_ and *T*_01_ < *T*_02_. Inspection of Eq. 4 and numerical simulation (Fig. 4A) show that if *p*_1_*T*_01_ = *p*_2_*T*_02_ (the case where *p*_1_ = 0.49, *p*_2_ = 0.21 TCID_50_cell^−1^d^−1^), then for fixed *β*_1_ + *β*_2_ *R*_0_ is independent of the individual values of *β*_1_ and *β*_2_. However, if *p*_1_*T*_01_ > *p*_2_*T*_02_ (the case where *p*_1_ = 0.63, *p*_2_ = 0.07 TCID_50_cell^−1^d^−1^), then *R*_0_ will increase with increasing *β*_1_ (and decreasing *β*_2_ to ensure *β*_1_ + *β*_2_ is fixed), and vice versa. Results when considering the AUC are qualitatively similar, although there are clear (but only quantitative) differences due to the non-linear dynamics of the full viral dynamics system which affect the AUC but not the calculation of *R*_0_.

**Fig. 4:**
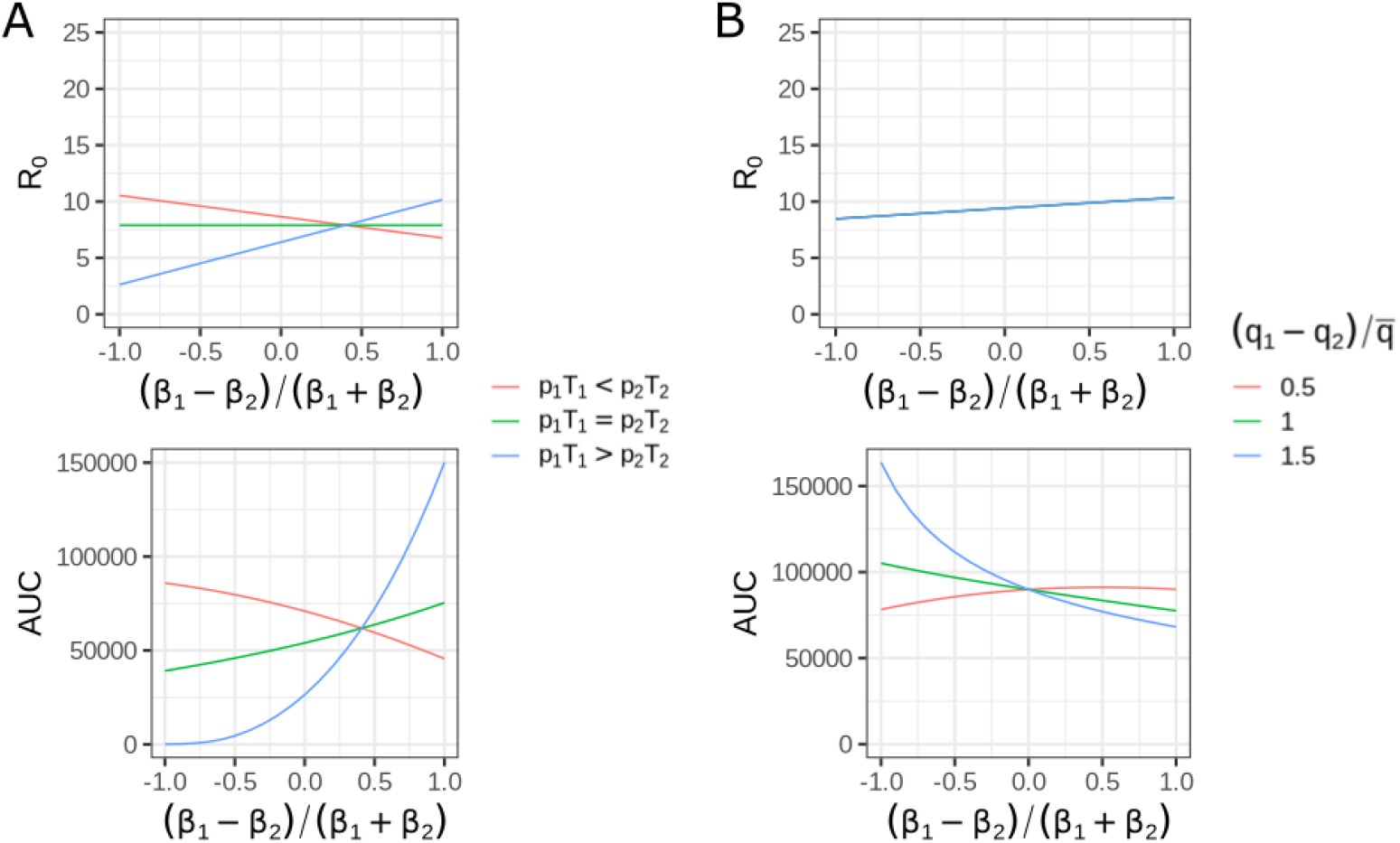
(Top) basic reproduction number and (bottom) viral loads for scenarios A, B and C when *β*_1_ − *β*_2_ is changed (x-axis) while *β*_1_ + *β*_2_ is held constant.

Then, we use the model to predict how viral fitness changes with cell tropism when the cell type with a higher viral production rate also has a higher interferon production rate (Fig. 4B). We find that there are three possible behaviours for the change in AUC (as a proxy for transmissibility) with respect to cell tropism (bottom panel). The first is that the increase in virus production for cell type 1 outweighs the increase in interferon production, so preferentially infecting cell type 1 increases transmissibility. For example, this happens for *p*_1_ = 0.385, *p*_2_ = 0.315 TCID_50_cell^−1^d^−1^, *q*_1_ = 1.25 × 10^−4^, *q*_2_ = 7.5 × 10^−5^ (red line in Fig. 4B). The second is that the increase in interferon production for cell type 1 outweighs the increase in interferon production, so preferentially infecting cell type 2 increases transmissibility. This happens for *p*_1_ = 0.385, *p*_2_ = 0.315 TCID_50_cell^−1^d^−1^, *q*_1_ = 1.5 × 10^−4^, *q*_2_ = 5 × 10^−5^ (green line in Fig. 4B). The last is that transmissibility changes non-monotonically with respect to cell type preference, and is highest for an intermediate value of cell tropism. This happens for *p*_1_ = 0.385, *p*_2_ = 0.315 TCID_50_cell^−1^d^−1^, *q*_1_ = 1.75 *×* 10^−4^, *q*_2_ = 2.5 *×* 10^−5^ (blue line in Fig. 4B). However, in all three of these cases, the within-host *R*_0_ increases with an increased preference for cell type 1, as changing *q*_*i*_ does not affect *R*_0_.

### 3.3 For models lacking innate and adaptive immunity, changes in transmissibility due to changes in cell tropism are sensitive to the choice of transmissibility metric

Previous studies have used different functions to map between the viral load and transmissibility (see comparison in Asher et al. (2023)). We repeated the analysis in Scenario A of Section 3.1, but with two alternative functions for the mapping instead of the area under the viral load curve (AUC) in linear space 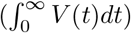 which was originally used. The first is the area under the viral load curve in log space, 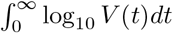. The second is the area under a dose–response transform of the viral load where transmissibility saturates with increasing viral load, 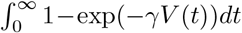, using *γ* = 10^−4^. The top row of Fig. 5 shows the values of each metric as *β*_1_ − *β*_2_ is varied, and cell type 1 is more abundant (the top left figure is the same as the bottom left figure of Fig. 3). Monotonicity is maintained regardless of the metric chosen: each metric increased as *β*_1_ − *β*_2_ increased.

**Fig. 5:**
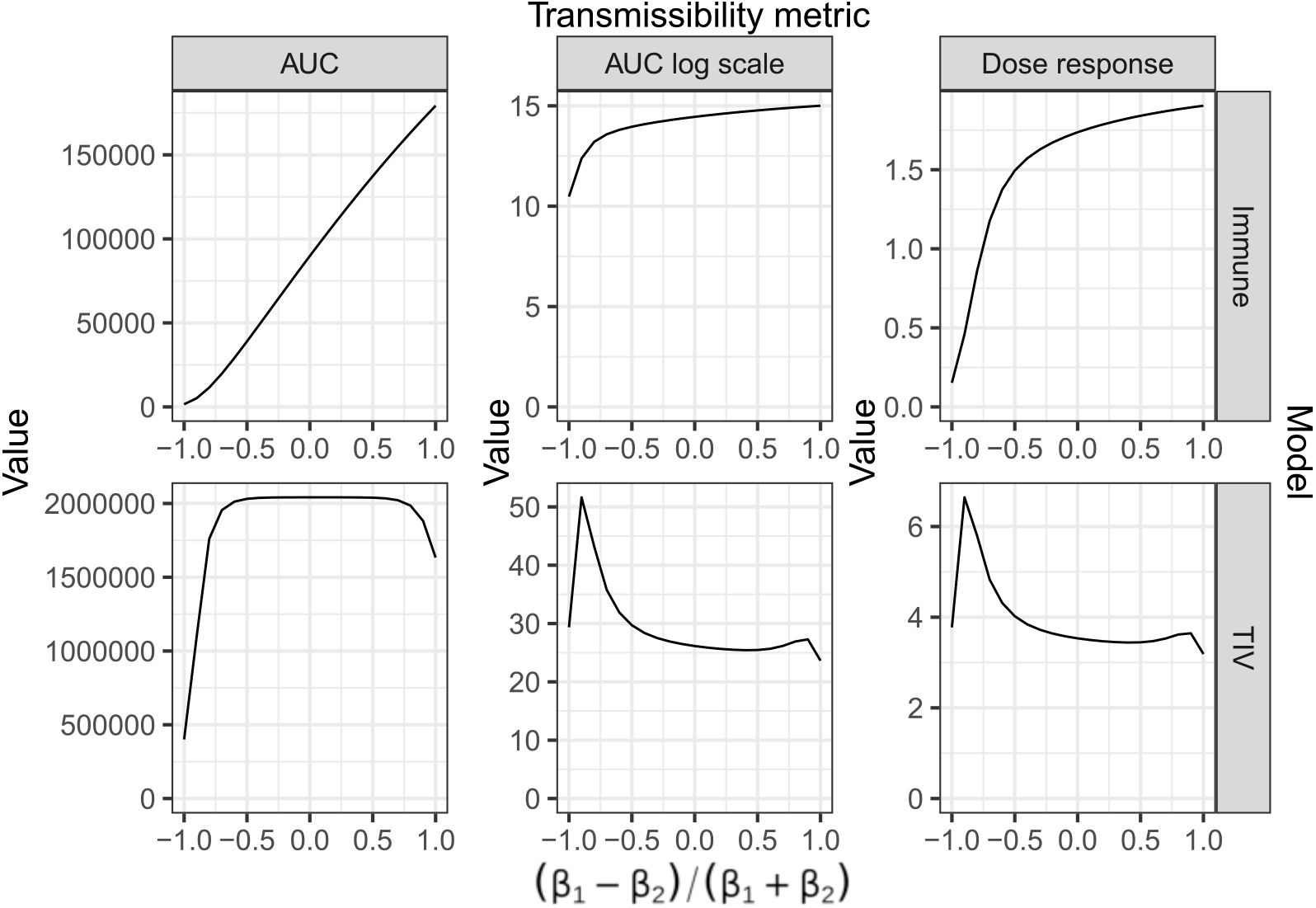
(*β*_1_ *β*_2_)/(*β*_1_ + *β*_2_) (x-axis) vs. values of transmissibility metrics (y-axis), for (top row) a model with innate and adaptive immunity, and (bottom row) the TIV model. The metrics are (from left to right) 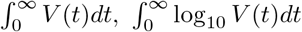, and 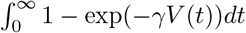, using *γ* = 10^−4^.

We repeated the analysis examining three different forms for the relationship between the viral load and transmissibility, but for a model with no innate and adaptive immunity, i.e. the TIV model found commonly in the viral dynamics literature (Baccam et al. 2006). We reduced the model in Eq. 2 to the TIV model by setting *A*(0) = 0 and *q*_*i*_ = 0. For this model, the AUC in linear space mostly increases as *β*_1_ − *β*_2_ increases, while the AUC in log space and the area under the dose-response transformed viral load curve mostly decrease (with some non-monotonicity towards extreme values of *β*_1_ − *β*_2_).

Accordingly, we plot the viral load trajectories for the TIV model (Fig. 6). As *β*_1_ − *β*_2_ increases, the initial growth rate increases, leading to a faster, higher peak time, but also a much faster resolution of the infection. In the TIV model, resolution of the infection is driven by target cell depletion, so a higher initial viral growth rate leads to faster target cell depletion and a faster resolution of the infection. Comparing the three metrics, the AUC in log space and the area under the dose-response transformed viral load curve are relatively more heavily influenced by the duration of the infection whereas the AUC in linear space is more heavily influenced by the peak viral load. Hence, the metrics’ changes in response to a viral load with a higher peak titre and shorter duration are different. For the model with innate and adaptive immune responses, the resolution of infection is due to the adaptive immune response and not target cell depletion, so the duration of infection is less sensitive to changes in cell susceptibility (Fig. 2A). Thus, increases in *β*_1_ − *β*_2_ increase all metrics.

**Fig. 6:**
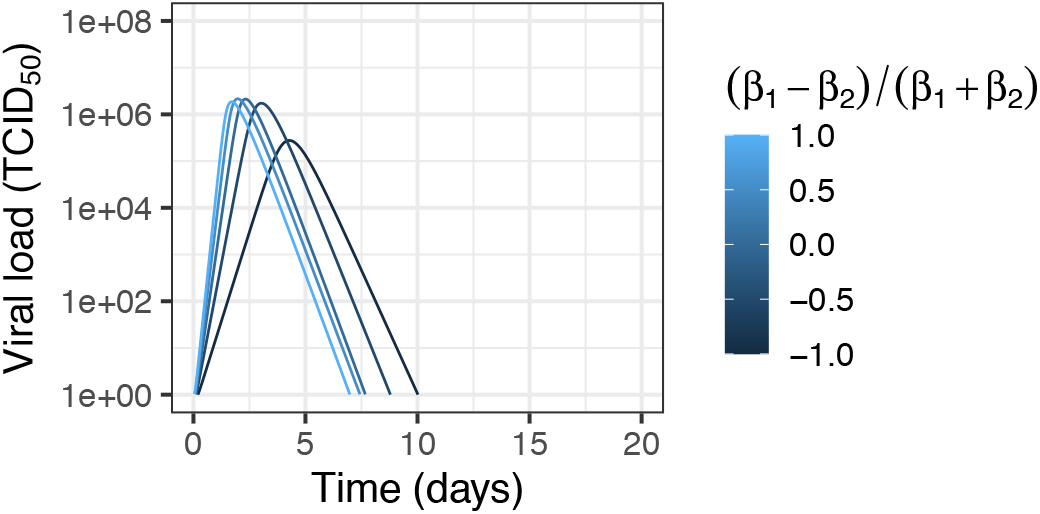
Viral load curves for the TIV model when 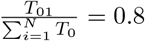, as *β*_1_ −*β*_2_ is changed while *β*_1_ + *β*_2_ is held constant.

### 3.4 Parameters which were heterogeneous between compartments could be estimated with the right type of data

So far, we have used models to qualitatively predict how changing cell tropism changes viral fitness. However, to use the models to make quantitative predictions about viruses of biological interest using experimental data, we need to be able to estimate model parameters, in particular the amount of heterogeneity between cells.

To investigate whether the values of model parameters can be estimated and used to make useful predictions, we conducted a simulation estimation study. We used the model with heterogeneity in *β*_*i*_ and *T*_0*i*_ only, and *N* = 2 as an example. We estimated *β*_1_ and *β*_2_ only. In practice, many other parameters for viral dynamics models are likely to be unidentifiable from available data (Tuncer et al. 2025), requiring alternative parameterisation strategies such as constraining parameters to values found in the literature. Here we focus on the identifiability of the parameters which are heterogeneous between cell types, and thus assume that all other parameters are known. We assume that *T*_0*i*_ were known as the initial distribution of cell types is readily measurable using common techniques such as flow cytometry.

We reparameterised *β*_1_ and *β*_2_ in terms of the weighted mean 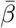 and a parameter summarising heterogeneity in susceptibility between cell types, Δ (Methods). We simulated 100 sets of parameters varying 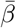, *δ, p/δ, c, V*_0_ and *T*_01_*/T*_0_ (Methods). For each parameter set we simulated data using Δ = .8, Δ = 0 and Δ = −0.8 (heterogeneity is maximal when Δ = ±1; it is difficult to establish reasonable values of Δ based on prior knowledge, as detailed in the Discussion). We then re-estimated 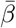 and Δ from either viral load data alone, or data on the proportion of uninfected and infected cells of each type over time in addition to viral load data.

Δ can be estimated with both viral load and cell proportion data. Fig. 7A shows the median (dots) and 95% credible intervals (error bars) for Δ for different true values of 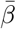 (x-axis) and Δ (top captions), when 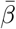 and Δ were inferred with viral load and cell proportion data. When the true value of Δ was −0.8, heterogeneity could be detected 83% of the time (that is, 83% of simulation–estimation studies recovered 95% credible intervals (CIs) for Δ which excluded 0). When the true value of Δ was −0.8, heterogeneity could be detected 75% of the time. We also tested whether there were false positives, i.e., heterogeneity was inferred even when the true value of Δ was zero. Only 3.5% of simulation–estimation studies where the true value of Δ was zero had 95% CIs for Δ excluding zero, so the rate of false positives was low. When we repeated the inference with viral load data only (Fig. 7B), the credible intervals spanned the prior distribution for Δ (uniform between −1 and 1), showing that Δ was not identifiable.

**Fig. 7:**
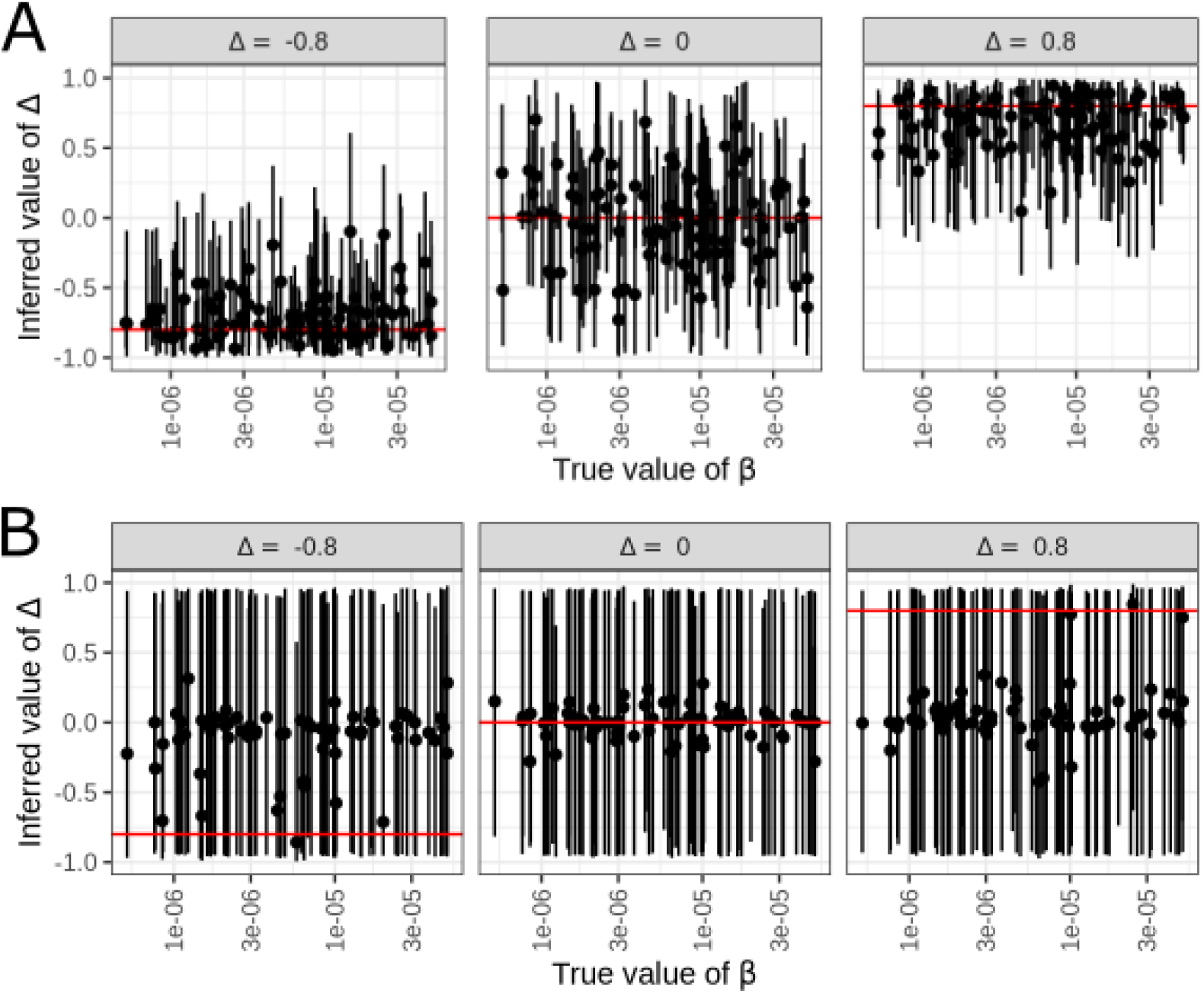
Median (dots) and 95% credible intervals (error bars) for Δ for different true values of 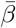 (x-axis) and Δ (top captions). Values in (A) were inferred with viral load and cell proportion data; values in (B) were inferred with viral load data only.

The result that data on cell proportion improve inference of Δ makes sense because if we have measurements of the relative numbers of infected cells in each compartment, these numbers reflect the relative susceptibility of these cell types of infection, whereas from the total viral load, the proportions of virus produced by each cell type are unknown.

Where individual model parameters are not identifiable (by examining the marginal posterior distribution), the joint posterior distribution may nonetheless be constrained enough to make precise model predictions. These predictions may be more important than the identifiability of model parameters themselves, for example if the main purpose of the study is to predict the impact of pharmaceutical interventions. We thus tested whether the difference in the identifiability of Δ with or without cell proportion data impacts the model’s ability to predict the viral load for hypothetical anatomical sites of infection with different initial proportions of cell types. To make these model predictions, we sampled from the joint posterior distribution of 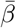 and Δ and used Eq. 8 to calculate *β*_*i*_. Then, holding these values of *β*_*i*_ constant, we varied *T*_0*i*_ while keeping their sum constant, and solved Eq. 2 to predict the viral load.

Figure 8 shows the predictions, according to the true and fitted model parameters, for how the viral load changes as the initial cell type proportion is changed. In Fig. 8A, the lines show the model predictions for the viral load using parameter set 1, varying the initial proportion of cell type 1 (colours) and Δ (columns). When Δ = 0, the true susceptibilities of the two cell types are equal, so the true model parameters predict no change in viral load regardless of the initial cell type proportion. When Δ = −0.8, then the susceptibility of cell type 1 to infection is higher than that of cell type 2 (*β*_1_ > *β*_2_). Therefore, increasing the initial proportion of cell type 1 increases the initial viral growth rate and peak viral load according to the true model parameters (left column of Fig. 8A). The opposite is true when Δ = 0.8. In this case, the susceptibility of cell type 2 to infection is higher than that of cell type 1 (*β*_2_ > *β*_1_). Therefore, increasing the initial proportion of cell type 1 decreases the initial viral growth rate and peak viral load according to the true model parameters (right column of Fig. 8A).

**Fig. 8:**
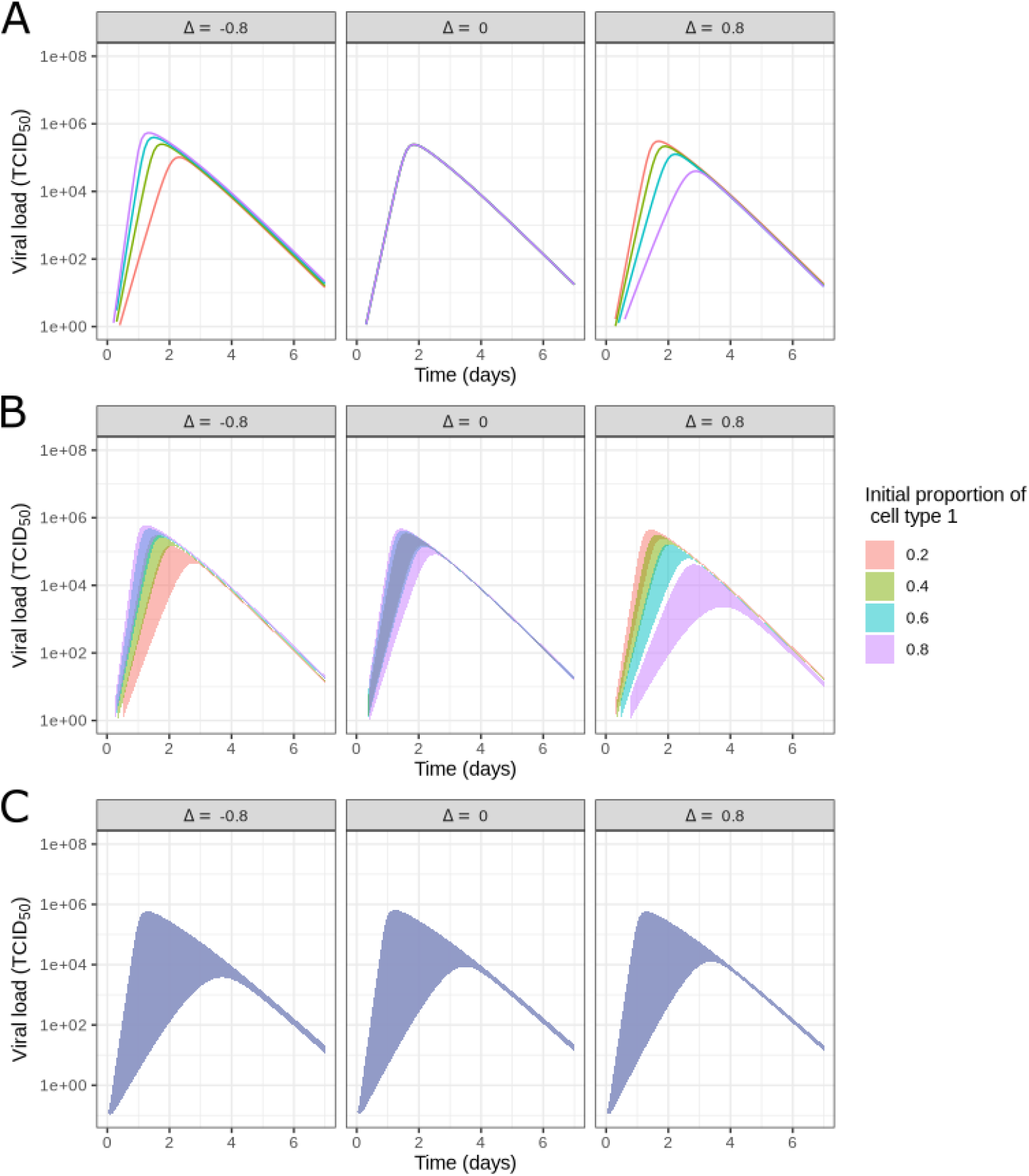
(A) Model predictions for the viral load using parameter set 1, varying the initial proportion of cell type 1 (colours) and Δ (columns) while keeping *β*_*i*_ and all other parameters constant at their true values. (B-C) 95% credible intervals for model predictions using samples from the posterior distribution for *β*_*i*_ rather than using the true values for *β*_*i*_. Predictions in (B) were made using the posterior distribution inferred with viral load and cell proportion data; predictions in (C) were made using the posterior distribution inferred with viral load data only.

Fig. 8B shows model predictions for the same initial cell type proportions as Fig. 8A, but made using parameter sets sampled from the posterior distribution, when viral load and cell proportion data (generated using the true values of Δ in the column labels) were used for model fitting. The shaded areas show the 95% credible intervals. The model fitted to the data generated assuming no cell type preference (Δ = 0) correctly predicts that the viral load is independent of the initial cell type proportions. It also correctly predicts from the data generated with a preference for cell type 1 (Δ = −0.8) that the growth rate and peak viral load would increase with higher proportions of cell type 1 in this situation, and vice versa for the data generated with a preference for cell type 2 (Δ = 0.8). Fig. 8C shows predictions made using the posterior distribution, when only viral load data was used for fitting. The 95% credible intervals are wide and overlap for all values of *T*_01_, so the fitted model was unable to predict the effect of changing the initial proportion of cell types.

To see whether these results are representative of the many simulated data sets used to re-estimate model parameters, Figures 1–10 in the Supplementary Material show these results for 10 of the 100 simulated data sets, and for individual parameter sets sampled from the posterior distribution rather than the 95% credible interval. Generally, in the cases where the 95% CI was able to infer heterogeneity in *β*_*i*_, model predictions from individual samples reflected trends shown in the true values, while in the cases where the 95% CI was not able to infer heterogeneity in *β*_*i*_, model predictions using the posterior distribution were less accurate.

## 4 Discussion

In this study, we constructed mathematical models for acute viral infection within the host which incorporated between-cell heterogeneity in susceptibility to infection, rates of virus production, and/or rates of interferon production. We have shown that when there was heterogeneity in rates of virus production and/or rates of interferon production, then a virus’ fitness changes depending on its cell tropism, i.e. which cell types are preferentially infected. In the case where there is a single source of heterogeneity, viral fitness is highest when the virus strongly preferentially infected the cell type that has the highest level of virus production/lowest level of interferon production. In the case where there are multiple competing sources of heterogeneity, the cell tropism for which the fitness is highest depends on the balance between these competing sources.

The within-host basic reproduction number only changes as heterogeneity is introduced in cells’ susceptibility to infection, rates of virus production and initial abundance, and does not change when rates relating to the innate immune response are changed. Moreover, where there is heterogeneity in cells’ rates of virus production and/or initial abundance, the basic reproduction number changes linearly with respect to changes in the amount of heterogeneity in cell types’ susceptibility to infection. On the other hand, when considering the virus’ ability to transmit to another host, we use total virus shedding as the fitness metric. This metric is affected by the innate immune response in addition to susceptibility and virus production. It can also change non-monotonically when there are multiple sources of heterogeneity, such that the highest fitness results from an intermediate difference between susceptibilities of cell types.

We tested three mathematical forms for the relationship between the the viral load curve and transmissibility, and found that model predictions of the change in transmissibility when heterogeneity in cells’ susceptibility to infection changed were qualitatively similar regardless of the mathematical form used. However, when the TIV model was used instead of a model with innate and adaptive immunity, then two of the three metrics predicted that a stronger preference for infecting more abundant cells, while increasing the within-host basic reproduction number, would ultimately decrease transmissibility through reducing the duration of infection. A similar result holds true for the TIV model even in the absence of heterogeneity: increasing *β* and/or *p* increases the within-host basic reproduction number while decreasing the duration of infection. To the best of the authors’ knowledge, there is no evidence for such a negative correlation between a virus’ within-host basic reproduction number and its duration of infection, or between a virus’ within-host basic reproduction number and population-level reproduction number (a measure of transmissibility). Conversely, animal studies have shown a positive correlation between an early peak in viral titre (higher basic reproduction number) and transmissibility as measured by the number of infected contact animals (Kieran et al. 2024). Thus, while the TIV model is useful for many contexts such as estimating a viral load curve with limited measurements, predictions made by varying its model parameters may not be accurate even at a qualitative level.

Two previous studies have constructed mathematical models of infection with two target cell populations. In the first of these studies (Dobrovolny et al. 2010), the parameter values for the primary cell type were held constant, while the parameter values for the secondary cell type were changed. However, when a virus evolves to infect different cell types, it is more likely that its characteristics (and so modelled parameter values) for both cell types will change simultaneously. In the second study (Reperant et al. 2012), the model was highly detailed, with compartments and parameterisations specifically for the influenza context. The study did not focus on the general qualitative behaviour of models featuring multiple target cell populations. While these models addressed the specific research questions posed by those studies, our analysis tackles the knowledge gaps of (a) how different sources of heterogeneity between cells affect within-host fitness and transmissibility, and (b) the feasibility of estimating parameters associated with cell tropism from data.

This study has a number of limitations. The greatest limitation is that in the simulations to explore model behaviour, it is difficult to know what realistic values are for the degree of cell tropism and other sources of heterogeneity. Previous studies of cell tropism have either treated tropism as a binary property, where cells either do or do not have the receptors necessary for virus binding (Hikmet et al. 2020), or have treated cell tropism as a qualitative property, noting a higher or lower proportion of infected cells for different cell types (Scull et al. 2009). As an alternative, flow cytometry or single-cell RNAseq data could be used to better quantify the proportion of each infected cells for each cell type. However, if there are sources of heterogeneity between cells other than unequal susceptibilities, then the proportions of infected cells may not directly correspond to differences in susceptibilities. Thus, mathematical models to estimate the differences between cell types would be required. In the simulation-estimation study, we used either equal susceptibilities to infection or a nine-fold difference in susceptibility (Δ = ±0.8). Once these models have been calibrated to data, then we could repeat the analysis with higher confidence. The result that the within-host basic reproduction number only changes as heterogeneity is introduced in cells’ susceptibility to infection, rates of virus production and initial abundance, and does not change when rates relating to the innate immune response are changed, is due to the model structure and is insensitive to the choice of parameter values. However, other results in the study could be sensitive to the choice of parameter values.

We focused on introducing heterogeneity to one baseline viral dynamics model for simplicity, but it is plausible that the relationship between between-cell heterogeneity and viral fitness could be qualitatively different for different model formulations. Mechanisms which were absent from our model include target cell natural death and (re)growth; production of non-infectious virus, such as defective interfering particles; and natural killer cells and CD8+ T cells. However, the formulae for the basic reproduction number and growth rate are insensitive to different models of the adaptive immune response. Mathematical models with more components are also more likely to have unidentifiable parameters when fitted to limited data. Thus, the realism of the baseline model must be balanced with the data available. The sensitivity of results to baseline parameter values could also be explored systematically in future work.

We did not consider the case where the infected cell lifetime varied between cell types. This was to keep the generation time constant regardless of heterogeneity. If the generation time remains constant as the amount of heterogeneity is changed, then there is a monotonic relationship between *R*_0_ and the growth rate *r*. Therefore, all results pertaining to *R*_0_ will also be true for *r*. However, if the infected cell lifetime varies between cell types, then introducing heterogeneity can vary the generation time, and results which are true for *R*_0_ may no longer be true for *r*. If this is the case, then more care has to be taken when comparing the fitnesses of viruses. For example, if one virus has a higher within-host *R*_0_ compared to another virus but a lower value of *r*, it is not immediately clear which virus will be fitter in the full life cycle, as the virus which replicates more quickly may gain an evolutionary advantage despite producing fewer secondary virions per generation.

This study captured heterogeneity between cells in terms of their “cell type” i.e. phenotypic characteristics. However, further heterogeneity could arise spatially. Spatial heterogeneity arises partly because of both short-range and long-range modes of spread for respiratory viruses (Zeng et al. 2022), rather than the system being well-mixed as assumed by the compartmental model approach. Further heterogeneity could arise if the cell types were not distributed randomly, but according to some spatial structure. For example, influenza A virus buds from the apical surface of cells (Boulan and Sabatini 1978), in proximity to goblet cells and ciliated cells but far from basal cells which are attached to the basement membrane. The effect of the spatial location of different cell types on viral fitness, in addition to intrinsic differences, could be explored in future work using a variety of methods for spatially structured models (Williams et al. 2025).

Moreover, heterogeneity could be present even within discrete cell types. Russell et al. (2018) showed that when identical cells from an immortalised cell line were infected with influenza virus, there was extreme heterogeneity in the amount of virus produced per cell. The effect of this within-cell-type heterogeneity could be explored with partial differential equation models.

Challenges remain in transforming data into formats which are suitable to fitting of mathematical models. For example, there is no single marker for resistant cells, so identification of these cells would require combining information from the expression of multiple genes or proteins.

## Acknowledgments

Ada W. C. Yan was funded by an Imperial College Research Fellowship and Wellcome Trust Collaborative Award (200187/Z/15/Z). Steven Riley was funded by a Wellcome Trust Investigator Award (UK, 200861/Z/16/Z). James M. McCaw was supported by an Australian Research Council Laureate Fellowship (FL240100126).

## Notes

### Competing Interest Statement

The authors have declared no competing interest.

## References

Asher, J., Lemenuel-Diot, A., Clay, M., Durham, D.P., Mier-y-Teran-Romero, L., Arguello, C.J., Jolivet, S., Wong, D.Y., Kuhlbusch, K., Clinch, B., Charoin, J.-E.: Novel modelling approaches to predict the role of antivirals in reducing influenza transmission. PLOS Computational Biology 19(1), 1010797 (2023) 10.1371/journal.pcbi.1010797. Publisher: Public Library of Science. Accessed 2025-03-27

Baccam, P., Beauchemin, C., Macken, C.A., Hayden, F.G., Perelson, A.S.: Kinetics of Influenza A Virus Infection in Humans. Journal of Virology 80(15), 7590–7599 (2006) 10.1128/jvi.01623-05. Publisher: American Society for Microbiology. Accessed 2025-12-12

Boulan, E.R., Sabatini, D.D.: Asymmetric budding of viruses in epithelial monlayers: a model system for study of epithelial polarity. Proceedings of the National Academy of Sciences 75(10), 5071–5075 (1978) 10.1073/pnas.75.10.5071. Publisher: Proceedings of the National Academy of Sciences. Accessed 2026-01-06

Bajaria, S.H., Webb, G., Cloyd, M., Kirschner, D.: Dynamics of Naive and Memory CD4+ T Lymphocytes in HIV-1 Disease Progression. JAIDS Journal of Acquired Immune Deficiency Syndromes 30(1), 41 (2002). Accessed 2025-10-08

Carnell, R.: Lhs: Latin Hypercube Samples, (2024). https://CRAN.R-project.org/package=lhs

Cao, P., Yan, A.W.C., Heffernan, J.M., Petrie, S., Moss, R.G., Carolan, L.A., Guarnaccia, T.A., Kelso, A., Barr, I.G., McVernon, J., Laurie, K.L., McCaw, J.M.: Innate Immunity and the Inter-exposure Interval Determine the Dynamics of Secondary Influenza Virus Infection and Explain Observed Viral Hierarchies. PLOS Computational Biology 11(8), 1004334 (2015) 10.1371/journal.pcbi.1004334. Publisher: Public Library of Science. Accessed 2026-02-26

Dobrovolny, H.M., Baron, M.J., Gieschke, R., Davies, B.E., Jumbe, N.L., Beauchemin, C.A.A.: Exploring Cell Tropism as a Possible Contributor to Influenza Infection Severity. PLOS ONE 5(11), 13811 (2010) 10.1371/journal.pone.0013811. Publisher: Public Library of Science. Accessed 2025-10-08

Dahari, H., Feliu, A., Garcia-Retortillo, M., Forns, X., Neumann, A.U.: Second hepatitis C replication compartment indicated by viral dynamics during liver transplantation. Journal of Hepatology 42(4), 491–498 (2005) 10.1016/j.jhep.2004.12.017. Accessed 2025-10-08

Diekmann, O., Heesterbeek, J.A.P., Roberts, M.G.: The construction of next-generation matrices for compartmental epidemic models. Journal of the Royal Society Interface 7(47), 873–885 (2010) 10.1098/rsif.2009.0386.Accessed 2025-12-23

Elaiw, A.M.: Global properties of a class of HIV models. Nonlinear Analysis: Real World Applications 11(4), 2253–2263 (2010) 10.1016/j.nonrwa.2009.07.001. Accessed 2025-10-08

FitzJohn, R.: Odin: ODE Generation and Integration, (2024). https://github.com/mrc-ide/odin

Fiege, J.K., Thiede, J.M., Nanda, H.A., Matchett, W.E., Moore, P.J., Montanari, N.R., Thielen, B.K., Daniel, J., Stanley, E., Hunter, R.C., Menachery, V.D., Shen, S.S., Bold, T.D., Langlois, R.A.: Single cell resolution of SARS-CoV-2 tropism, antiviral responses, and susceptibility to therapies in primary human airway epithelium. PLOS Pathogens 17(1), 1009292 (2021) 10.1371/journal.ppat.1009292. Publisher: Public Library of Science. Accessed 2025-04-17

Gabry, J., Češnovar, R., Johnson, A., Bronder, S.: Cmdstanr: R Interface to ‘Cmd-Stan’, (2025). https://mc-stan.org/cmdstanr/

Handel, A., Jr, I.M.L., Antia R.: Neuraminidase Inhibitor Resistance in Influenza: Assessing the Danger of Its Generation and Spread. PLOS Computational Biology 3(12), 240 (2007) 10.1371/journal.pcbi.0030240. Publisher: Public Library of Science. Accessed 2020-11-18

Higgins, D., Looker, J., Sunnucks, R., Carruthers, J., Finnie, T., Keeling, M.J., Hill, E.M.: Introducing a framework for within-host dynamics and mutations modelling of H5N1 influenza infection in humans. Journal of The Royal Society Interface 22(228), 20240910 (2025) 10.1098/rsif.2024.0910. Publisher: Royal Society. Accessed 2025-10-08

Hikmet, F., Méar, L., Edvinsson, A., Micke, P., Uhlén, M., Lindskog, C.: The protein expression profile of ACE2 in human tissues. Molecular Systems Biology 16(7), 209610 (2020) 10.15252/msb.20209610. Accessed 2026-04-22

Kelly, J.N., Laloli, L., V’kovski, P., Holwerda, M., Portmann, J., Thiel, V., Dijkman, R.: Comprehensive single cell analysis of pandemic influenza A virus infection in the human airways uncovers cell-type specific host transcriptional signatures relevant for disease progression and pathogenesis. Frontiers in Immunology 13 (2022). Accessed 2022-10-07

Kieran, T.J., Sun, X., Maines, T.R., Belser, J.A.: Optimal thresholds and key parameters for predicting influenza A virus transmission events in ferrets. npj Viruses 2(1), 64 (2024) 10.1038/s44298-024-00074-w. Publisher: Nature Publishing Group. Accessed 2025-12-12

Lipsitch, M., Barclay, W., Raman, R., Russell, C.J., Belser, J.A., Cobey, S., Kasson, P.M., Lloyd-Smith, J.O., Maurer-Stroh, S., Riley, S., Beauchemin, C.A., Bedford, T., Friedrich, T.C., Handel, A., Herfst, S., Murcia, P.R., Roche, B., Wilke, C.O., Russell, C.A.: Viral factors in influenza pandemic risk assessment. eLife 5, 18491 (2016) 10.7554/eLife.18491. Publisher: eLife Sciences Publications, Ltd. Accessed 2024-12-23

Long, J.S., Giotis, E.S., Moncorgé, O., Frise, R., Mistry, B., James, J., Morisson, M., Iqbal, M., Vignal, A., Skinner, M.A., Barclay, W.S.: Species difference in ANP32A underlies influenza A virus polymerase host restriction. Nature 529(7584), 101–104 (2016) 10.1038/nature16474. Publisher: Nature Publishing Group. Accessed 2025-12-23

Lindeboom, R.G.H., Worlock, K.B., Dratva, L.M., Yoshida, M., Scobie, D., Wagstaffe, H.R., Richardson, L., Wilbrey-Clark, A., Barnes, J.L., Kretschmer, L., Polanski, K., Allen-Hyttinen, J., Mehta, P., Sumanaweera, D., Boccacino, J.M., Sungnak, W., Elmentaite, R., Huang, N., Mamanova, L., Kapuge, R., Bolt, L., Prigmore, E., Killingley, B., Kalinova, M., Mayer, M., Boyers, A., Mann, A., Swadling, L., Woodall, M.N.J., Ellis, S., Smith, C.M., Teixeira, V.H., Janes, S.M., Chambers, R.C., Haniffa, M., Catchpole, A., Heyderman, R., Noursadeghi, M., Chain, B., Mayer, A., Meyer, K.B., Chiu, C., Nikolić, M.Z., Teichmann, S.A.: Human SARS-CoV-2 challenge uncovers local and systemic response dynamics. Nature 631(8019), 189–198 (2024) 10.1038/s41586-024-07575-x. Publisher: Nature Publishing Group. Accessed 2025-10-07

Miao, H., Hollenbaugh, J.A., Zand, M.S., Holden-Wiltse, J., Mosmann, T.R., Perel-son, A.S., Wu, H., Topham, D.J.: Quantifying the Early Immune Response and Adaptive Immune Response Kinetics in Mice Infected with Influenza A Virus. Journal of Virology 84(13), 6687–6698 (2010) 10.1128/jvi.00266-10. Publisher: American Society for Microbiology. Accessed 2026-02-26

Perelson, A.S., Essunger, P., Cao, Y., Vesanen, M., Hurley, A., Saksela, K., Markowitz, M., Ho, D.D.: Decay characteristics of HIV-1-infected compartments during combination therapy. Nature 387(6629), 188–191 (1997) 10.1038/387188a0. Publisher: Nature Publishing Group. Accessed 2025-10-08

Pawelek, K.A., Huynh, G.T., Quinlivan, M., Cullinane, A., Rong, L., Perelson, A.S.: Modeling Within-Host Dynamics of Influenza Virus Infection Including Immune Responses. PLOS Computational Biology 8(6), 1002588 (2012) 10.1371/journal.pcbi.1002588. Publisher: Public Library of Science. Accessed 2025-12-10

Payne, R.J., Nowak, M.A., Blumberg, B.S.: Analysis of a cellular model to account for the natural history of infection by the hepatitis B virus and its role in the development of primary hepatocellular carcinoma. Journal of Theoretical Biology 159(2), 215–240 (1992) 10.1016/s0022-5193(05)80703-9

R Core Team: R: A Language and Environment for Statistical Computing. R Foundation for Statistical Computing, Vienna, Austria (2024). https://www.R-project.org/

Reperant, L.A., Kuiken, T., Grenfell, B.T., Osterhaus, A.D.M.E., Dobson, A.P.: Linking Influenza Virus Tissue Tropism to Population-Level Reproductive Fitness. PLOS ONE 7(8), 43115 (2012) 10.1371/journal.pone.0043115. Publisher: Public Library of Science. Accessed 2025-10-08

Roach, S.N., Shepherd, F.K., Mickelson, C.K., Fiege, J.K., Thielen, B.K., Pross, L.M., Sanders, A.E., Mitchell, J.S., Robertson, M., Fife, B.T., Langlois, R.A.: Tropism for ciliated cells is the dominant driver of influenza viral burst size in the human airway. Proceedings of the National Academy of Sciences 121(31), 2320303121 (2024) 10.1073/pnas.2320303121. Publisher: Proceedings of the National Academy of Sciences. Accessed 2025-03-20

Russell, A.B., Trapnell, C., Bloom, J.D.: Extreme heterogeneity of influenza virus infection in single cells. eLife 7, 32303 (2018) 10.7554/eLife.32303. Publisher: eLife Sciences Publications, Ltd. Accessed 2025-04-17

Scull, M.A., Gillim-Ross, L., Santos, C., Roberts, K.L., Bordonali, E., Subbarao, K., Barclay, W.S., Pickles, R.J.: Avian Influenza Virus Glycoproteins Restrict Virus Replication and Spread through Human Airway Epithelium at Temperatures of the Proximal Airways. PLOS Pathogens 5(5), 1000424 (2009) 10.1371/journal.ppat.1000424. Publisher: Public Library of Science. Accessed 2026-04-22

Smith, A.P., Moquin, D.J., Bernhauerova, V., Smith, A.M.: Influenza Virus Infection Model With Density Dependence Supports Biphasic Viral Decay. Frontiers in Microbiology 9 (2018) 10.3389/fmicb.2018.01554. Publisher: Frontiers. Accessed 2025-10-27

Sikkema, L., Ramírez-Suástegui, C., Strobl, D.C., Gillett, T.E., Zappia, L., Madis-soon, E., Markov, N.S., Zaragosi, L.-E., Ji, Y., Ansari, M., Arguel, M.-J., Apperloo, L., Banchero, M., Bécavin, C., Berg, M., Chichelnitskiy, E., Chung, M.-i., Collin, A., Gay, A.C.A., Gote-Schniering, J., Hooshiar Kashani, B., Inecik, K., Jain, M., Kapellos, T.S., Kole, T.M., Leroy, S., Mayr, C.H., Oliver, A.J., Papen, M., Peter, L., Taylor, C.J., Walzthoeni, T., Xu, C., Bui, L.T., De Donno, C., Dony, L., Faiz, A., Guo, M., Gutierrez, A.J., Heumos, L., Huang, N., Ibarra, I.L., Jackson, N.D., Kadur Lakshminarasimha Murthy, P., Lotfollahi, M., Tabib, T., Talavera-L ópez, C., Travaglini, K.J., Wilbrey-Clark, A., Worlock, K.B., Yoshida, M., Berge, M., Bossé, Y., Desai, T.J., Eickelberg, O., Kaminski, N., Krasnow, M.A., Lafyatis, R., Nikolic, M.Z., Powell, J.E., Rajagopal, J., Rojas, M., Rozenblatt-Rosen, O., Sei-bold, M.A., Sheppard, D., Shepherd, D.P., Sin, D.D., Timens, W., Tsankov, A.M., Whitsett, J., Xu, Y., Banovich, N.E., Barbry, P., Duong, T.E., Falk, C.S., Meyer, K.B., Kropski, J.A., Pe’er, D., Schiller, H.B., Tata, P.R., Schultze, J.L., Teichmann, S.A., Misharin, A.V., Nawijn, M.C., Luecken, M.D., Theis, F.J.: An integrated cell atlas of the lung in health and disease. Nature Medicine 29(6), 1563–1577 (2023) 10.1038/s41591-023-02327-2. Publisher: Nature Publishing Group. Accessed 2026-03-18

{Stan Development Team}: Stan Reference Manual Version 2.37.0, (2025). https://mc-stan.org

Tuncer, N., Martcheva, M., Ciupe, S.M.: Structural and practical identifiability of within-host models of virus dynamics—A review. Current Opinion in Systems Biology 42, 100552 (2025) 10.1016/j.coisb.2025.100552. Accessed 2025-10-08

Woodall, M.N.J., Cujba, A.-M., Worlock, K.B., Case, K.-M., Masonou, T., Yoshida, M., Polanski, K., Huang, N., Lindeboom, R.G.H., Mamanova, L., Bolt, L., Richard-son, L., Cakir, B., Ellis, S., Palor, M., Burgoyne, T., Pinto, A., Moulding, D., McHugh, T.D., Saleh, A., Kilich, E., Mehta, P., O’Callaghan, C., Zhou, J., Barclay, W., De Coppi, P., Butler, C.R., Cortina-Borja, M., Vinette, H., Roy, S., Breuer, J., Chambers, R.C., Heywood, W.E., Mills, K., Hynds, R.E., Teichmann, S.A., Meyer, K.B., Nikoli ć, M.Z., Smith, C.M.: Age-specific nasal epithelial responses to SARS-CoV-2 infection. Nature Microbiology 9(5), 1293–1311 (2024) 10.1038/s41564-024-01658-1. Publisher: Nature Publishing Group. Accessed 2026-03-18

Williams, T., McCaw, J.M., Osborne, J.M.: Spatially structured models of viral dynamics: a scoping review. Microbiology and Molecular Biology Reviews 89(4), 00283–24 (2025) 10.1128/mmbr.00283-24. Publisher: American Society for Microbiology. Accessed 2026-01-06

Wang, X., Song, X., Tang, S., Rong, L.: Analysis of HIV models with multiple target cell populations and general nonlinear rates of viral infection and cell death. Mathematics and Computers in Simulation 124, 87–103 (2016) 10.1016/j.matcom.2015.11.011. Accessed 2025-10-08

Yoshida, M., Worlock, K.B., Huang, N., Lindeboom, R.G.H., Butler, C.R., Kumasaka, N., Dominguez Conde, C., Mamanova, L., Bolt, L., Richardson, L., Polanski, K., Madissoon, E., Barnes, J.L., Allen-Hyttinen, J., Kilich, E., Jones, B.C., Wilton, A., Wilbrey-Clark, A., Sungnak, W., Pett, J.P., Weller, J., Prigmore, E., Yung, H., Mehta, P., Saleh, A., Saigal, A., Chu, V., Cohen, J.M., Cane, C., Iordanidou, A., Shibuya, S., Reuschl, A.-K., Herczeg, I.T., Argento, A.C., Wunderink, R.G., Smith, S.B., Poor, T.A., Gao, C.A., Dematte, J.E., Reynolds, G., Haniffa, M., Bowyer, G.S., Coates, M., Clatworthy, M.R., Calero-Nieto, F.J., Göttgens, B., O’Callaghan, C., Sebire, N.J., Jolly, C., De Coppi, P., Smith, C.M., Misharin, A.V., Janes, S.M., Teichmann, S.A., Nikolić, M.Z., Meyer, K.B.: Local and systemic responses to SARS-CoV-2 infection in children and adults. Nature 602(7896), 321–327 (2022) 10.1038/s41586-021-04345-x. Publisher: Nature Publishing Group. Accessed 2026-03-18

Yan, A.W.C., Zaloumis, S.G., Simpson, J.A., McCaw, J.M.: Sequential infection experiments for quantifying innate and adaptive immunity during influenza infection. PLOS Computational Biology 15(1), 1006568 (2019) 10.1371/journal.pcbi.1006568. Publisher: Public Library of Science. Accessed 2024-12-23

Zeng, C., Evans, J.P., King, T., Zheng, Y.-M., Oltz, E.M., Whelan, S.P.J., Saif, L.J., Peeples, M.E., Liu, S.-L.: SARS-CoV-2 spreads through cell-to-cell transmission. Proceedings of the National Academy of Sciences 119(1), 2111400119 (2022) 10.1073/pnas.2111400119. Publisher: Proceedings of the National Academy of Sciences. Accessed 2025-10-09

