## Supplementary Figures for "Modelling between-cell heterogeneity in within-host influenza virus infection"

### Supplementary Material for Modelling between-cell heterogeneity in within-host influenza virus infection

Ada Yan

May 17, 2026

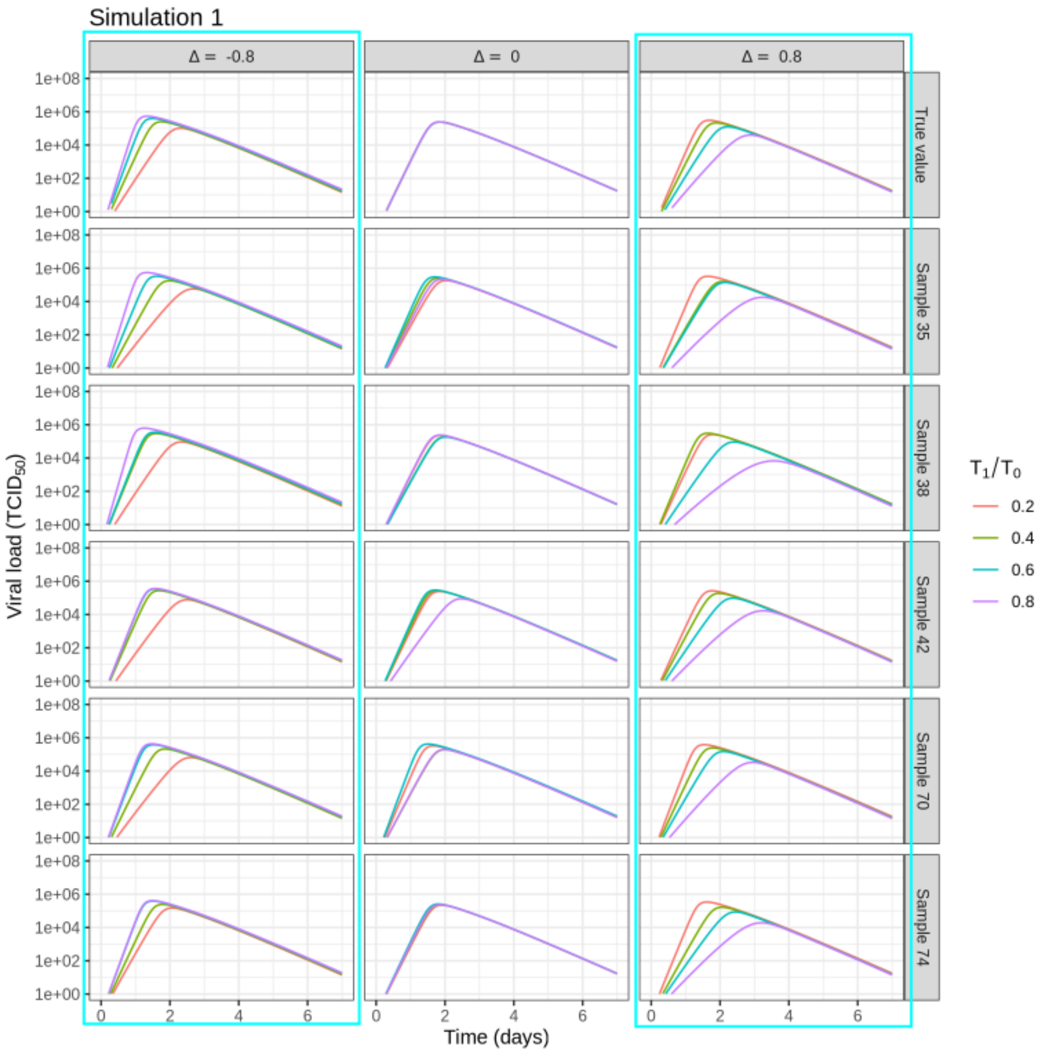


Figure 1: Model simulations with true and inferred parameter values for parameter set 1. Blue outlines indicate parameter sets where heterogeneity in $\beta_i$ was inferred.

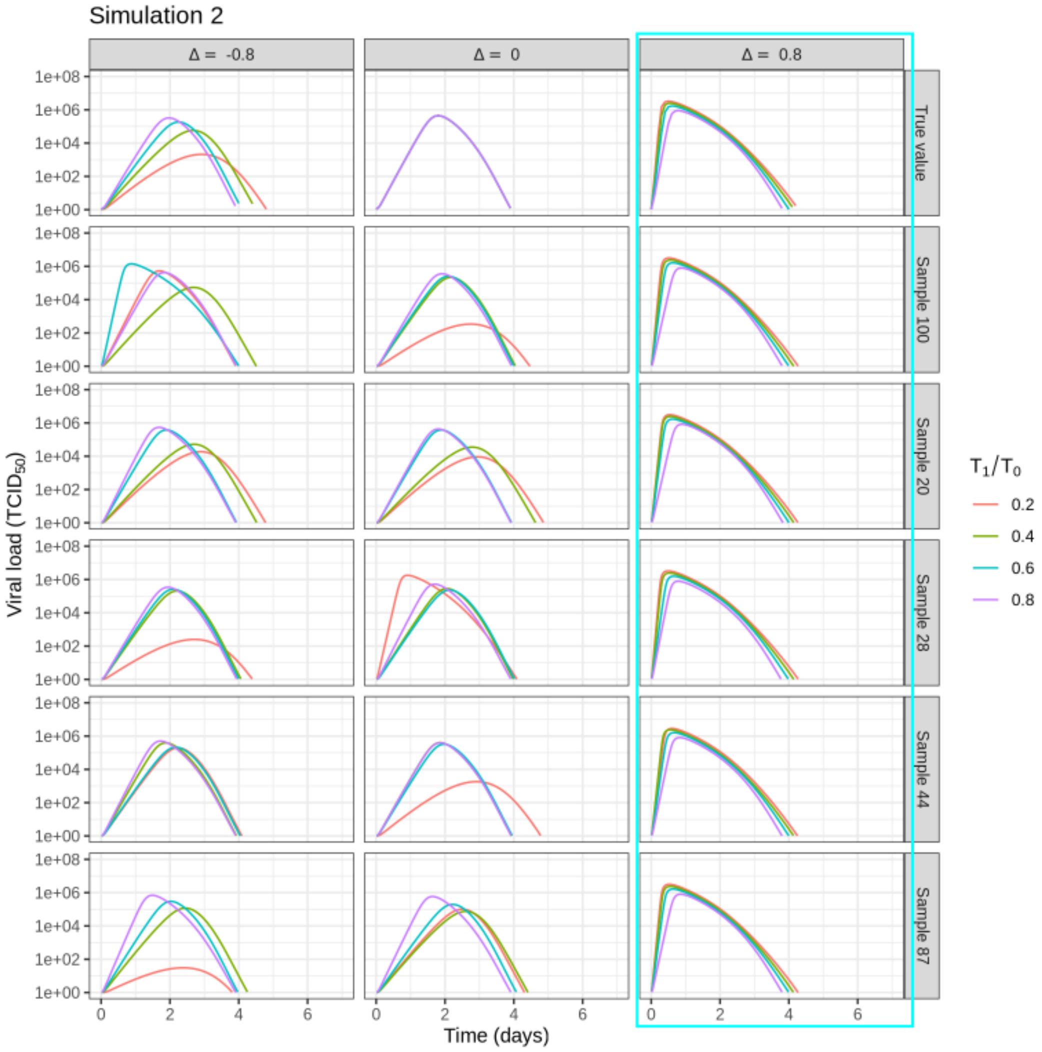


Figure 2: Model simulations with true and inferred parameter values for parameter set 2.

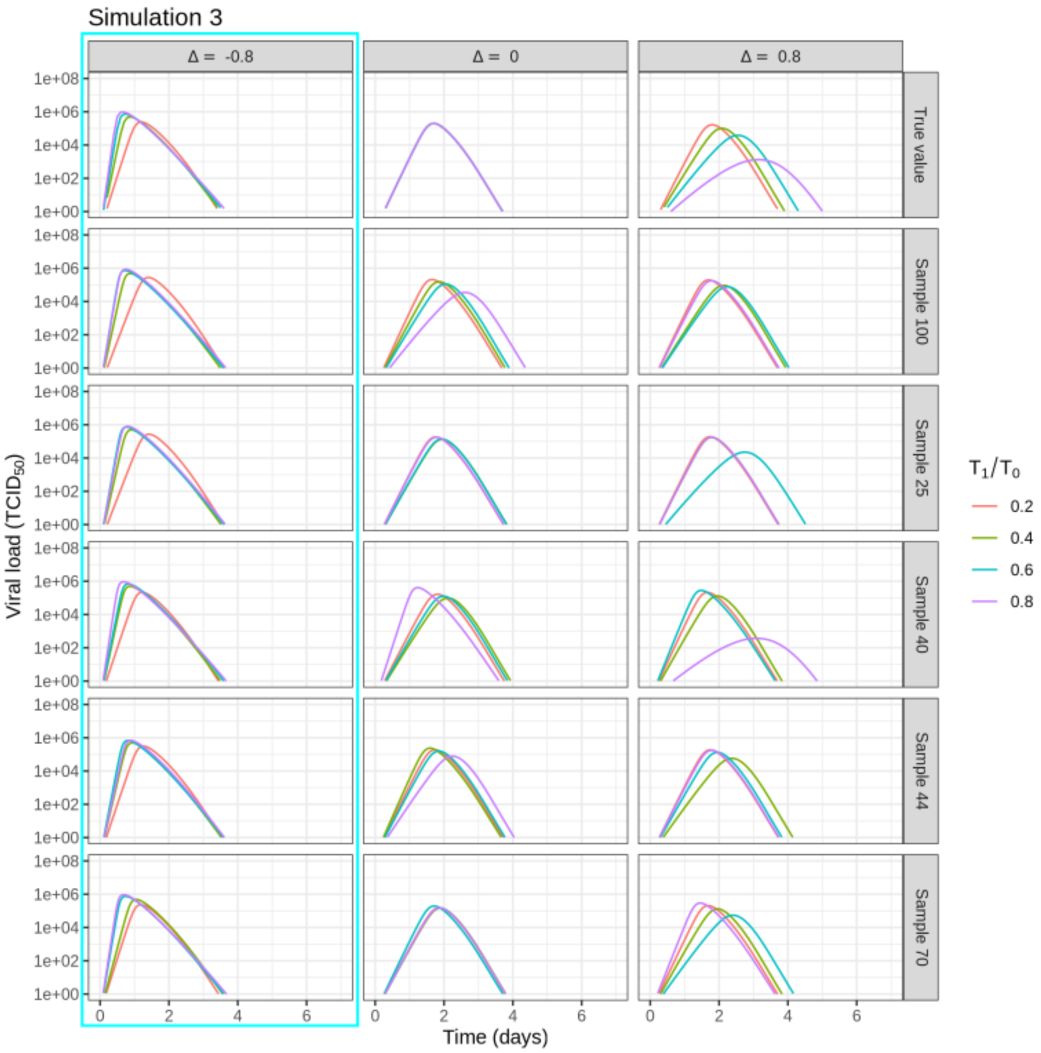


Figure 3: Model simulations with true and inferred parameter values for parameter set 3.

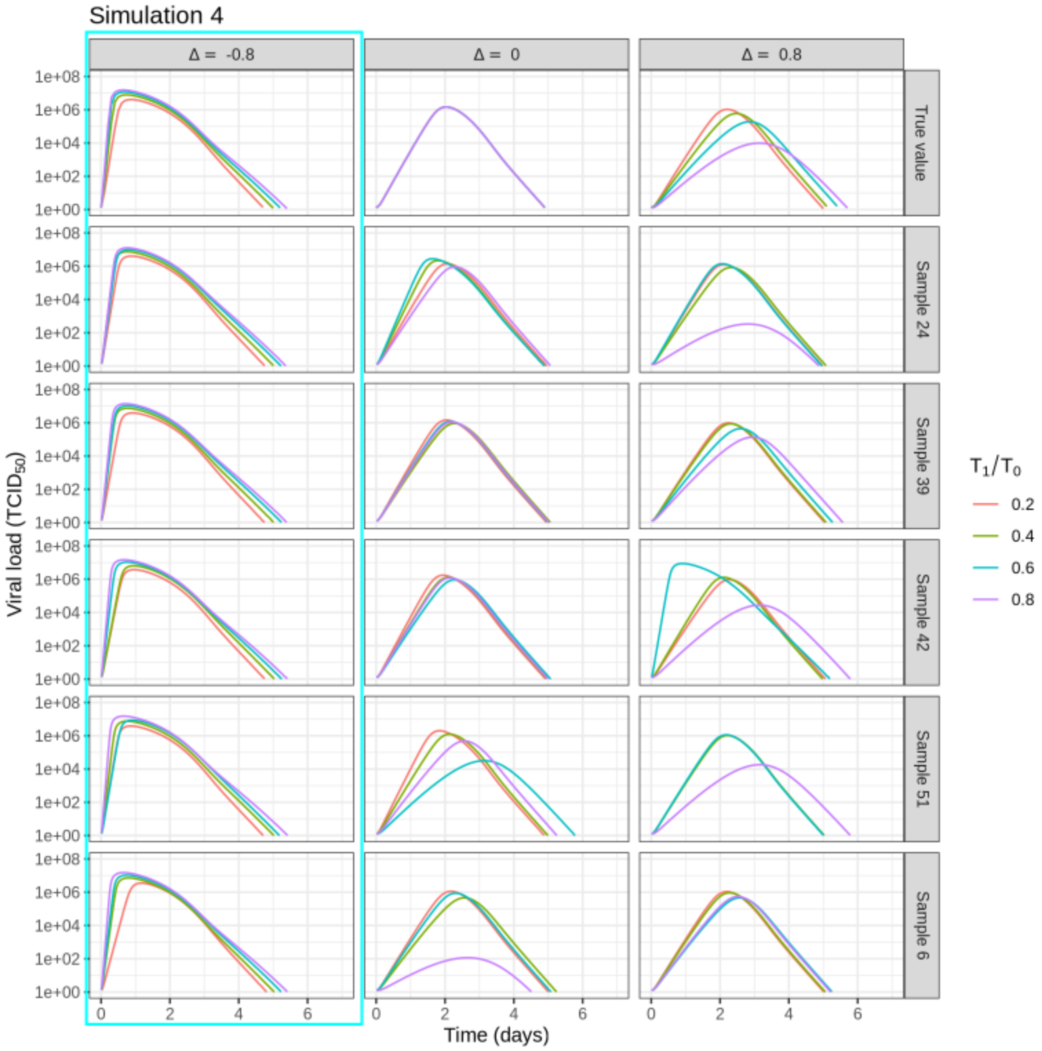


Figure 4: Model simulations with true and inferred parameter values for parameter set 4.

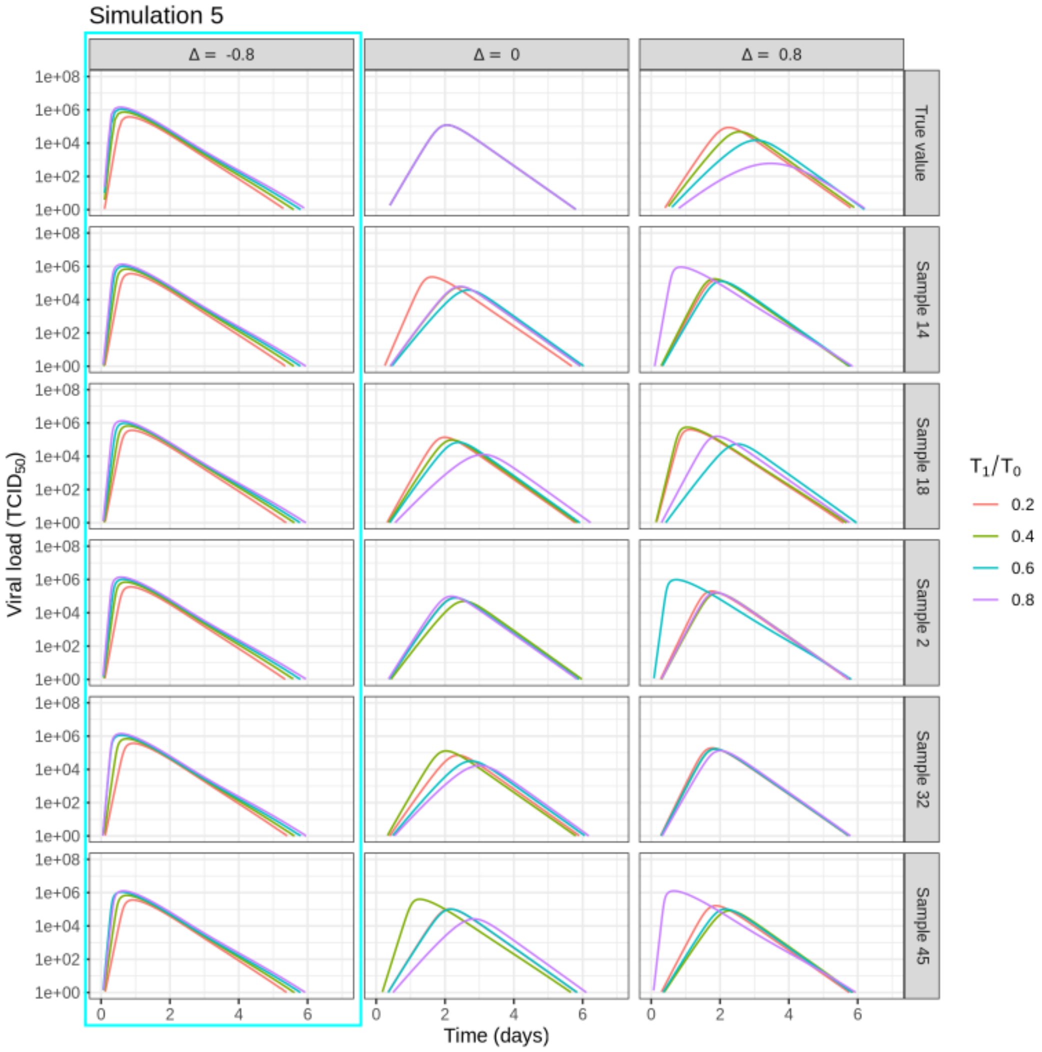


Figure 5: Model simulations with true and inferred parameter values for parameter set 5.

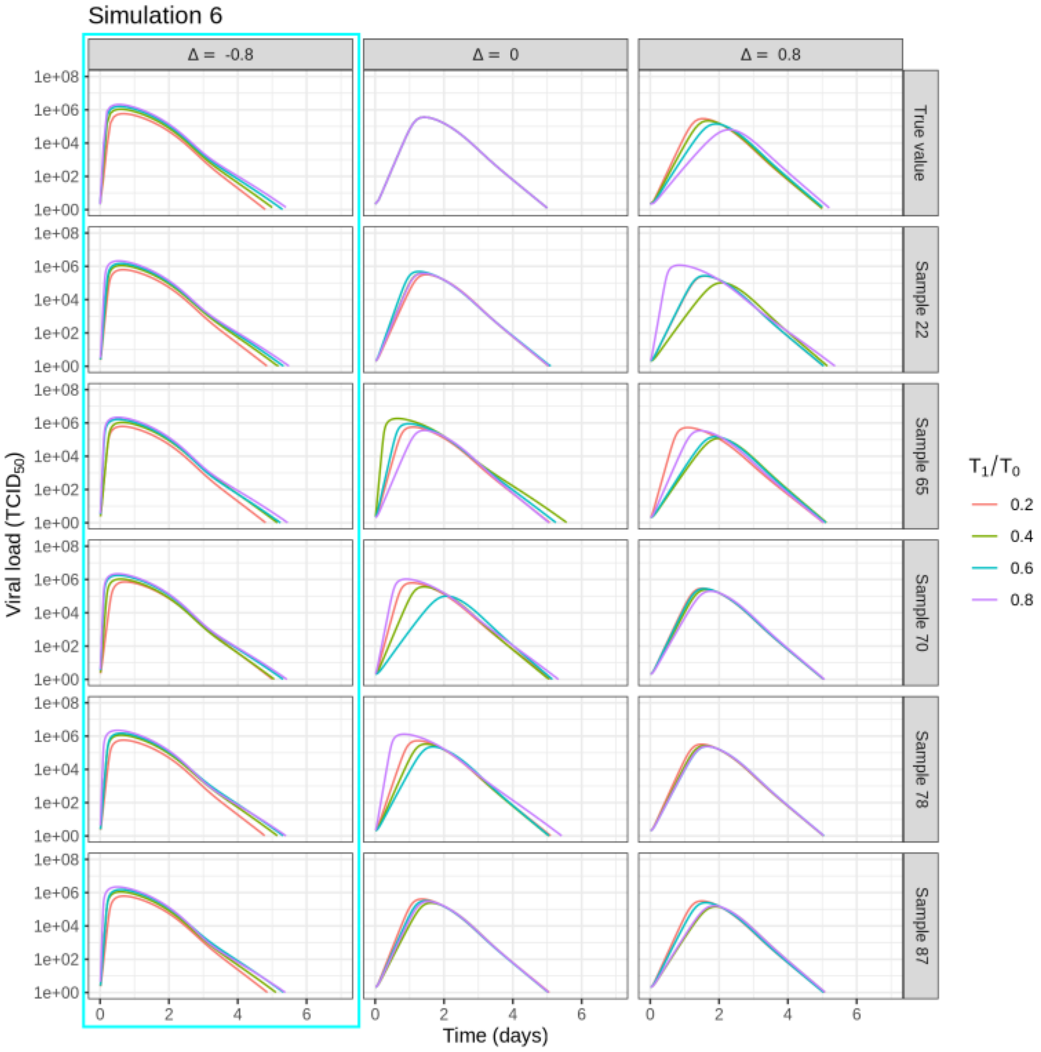


Figure 6: Model simulations with true and inferred parameter values for parameter set 6.

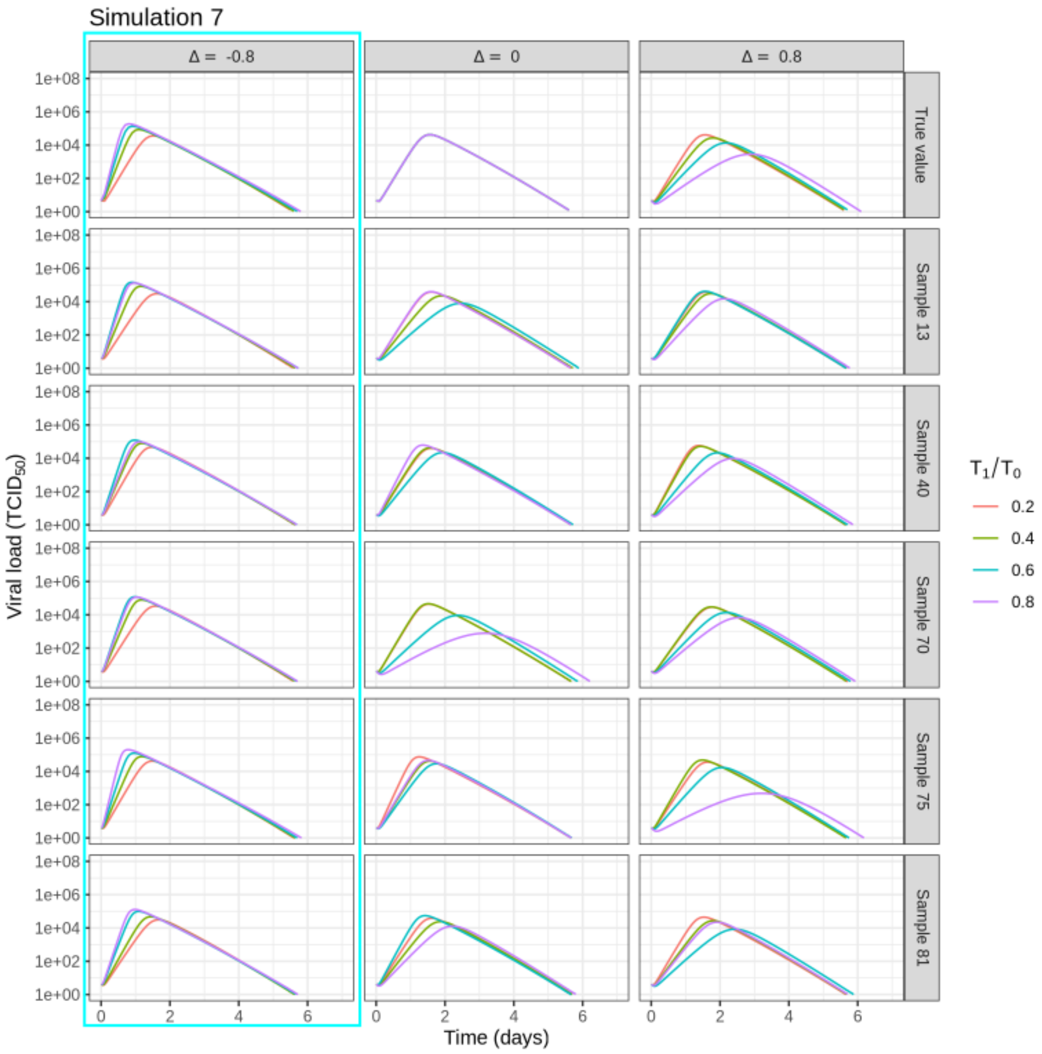


Figure 7: Model simulations with true and inferred parameter values for parameter set 7.

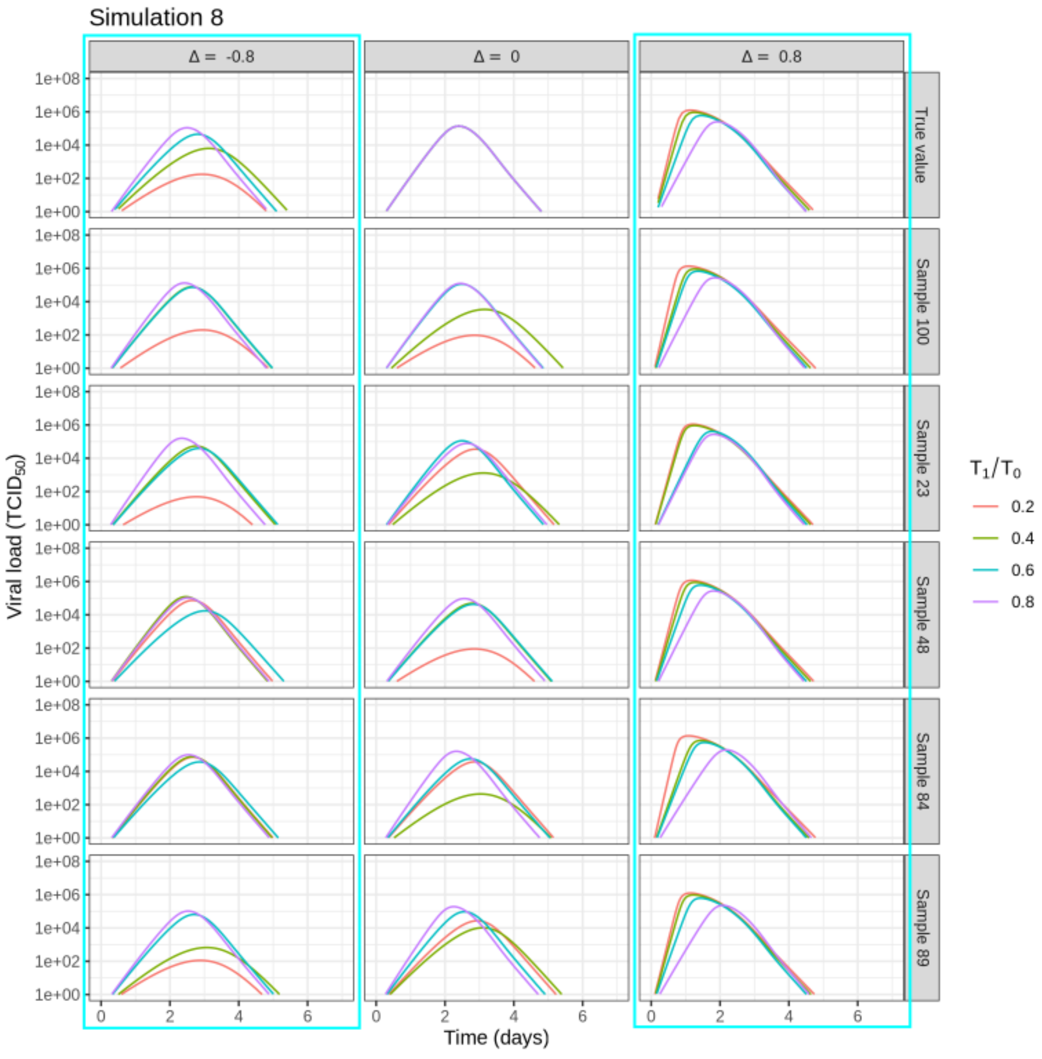


Figure 8: Model simulations with true and inferred parameter values for parameter set 8.

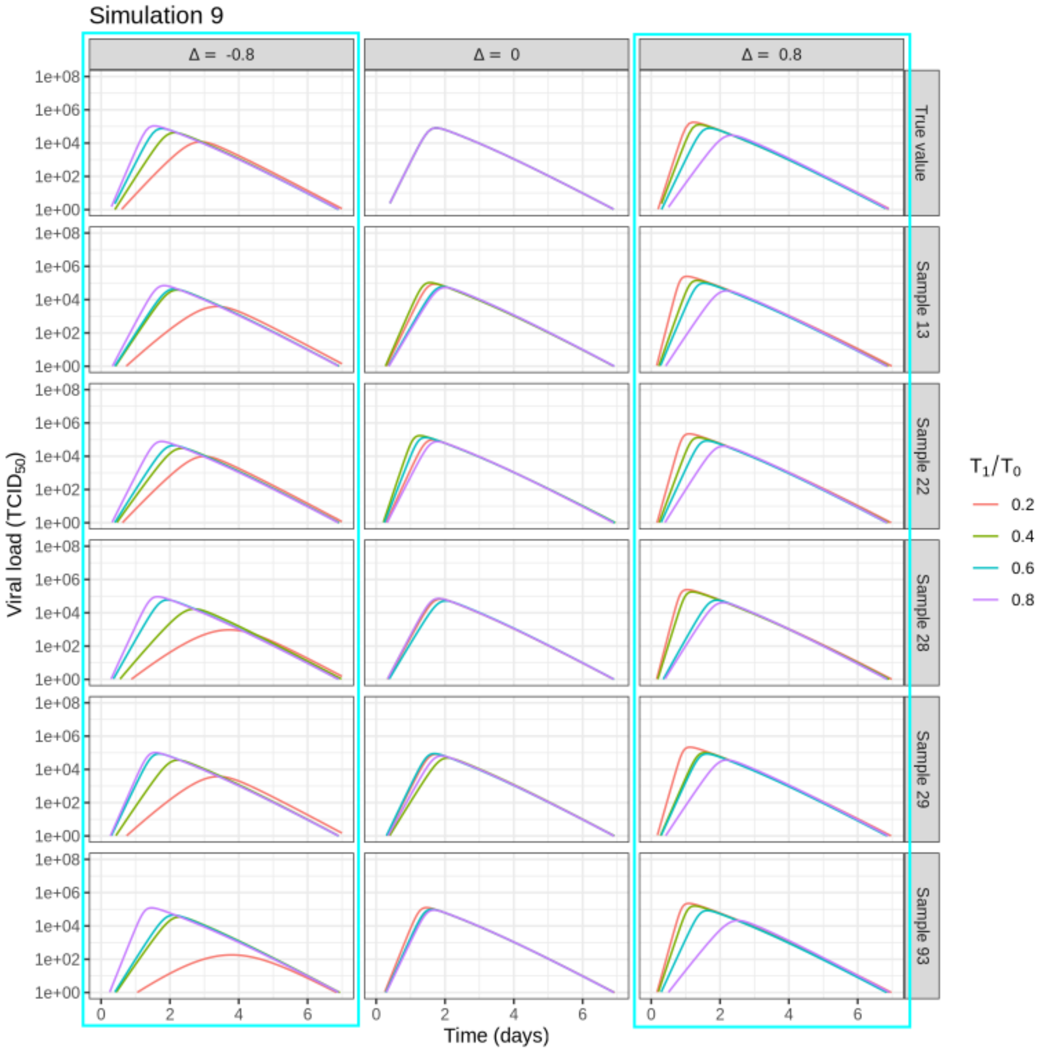


Figure 9: Model simulations with true and inferred parameter values for parameter set 9.

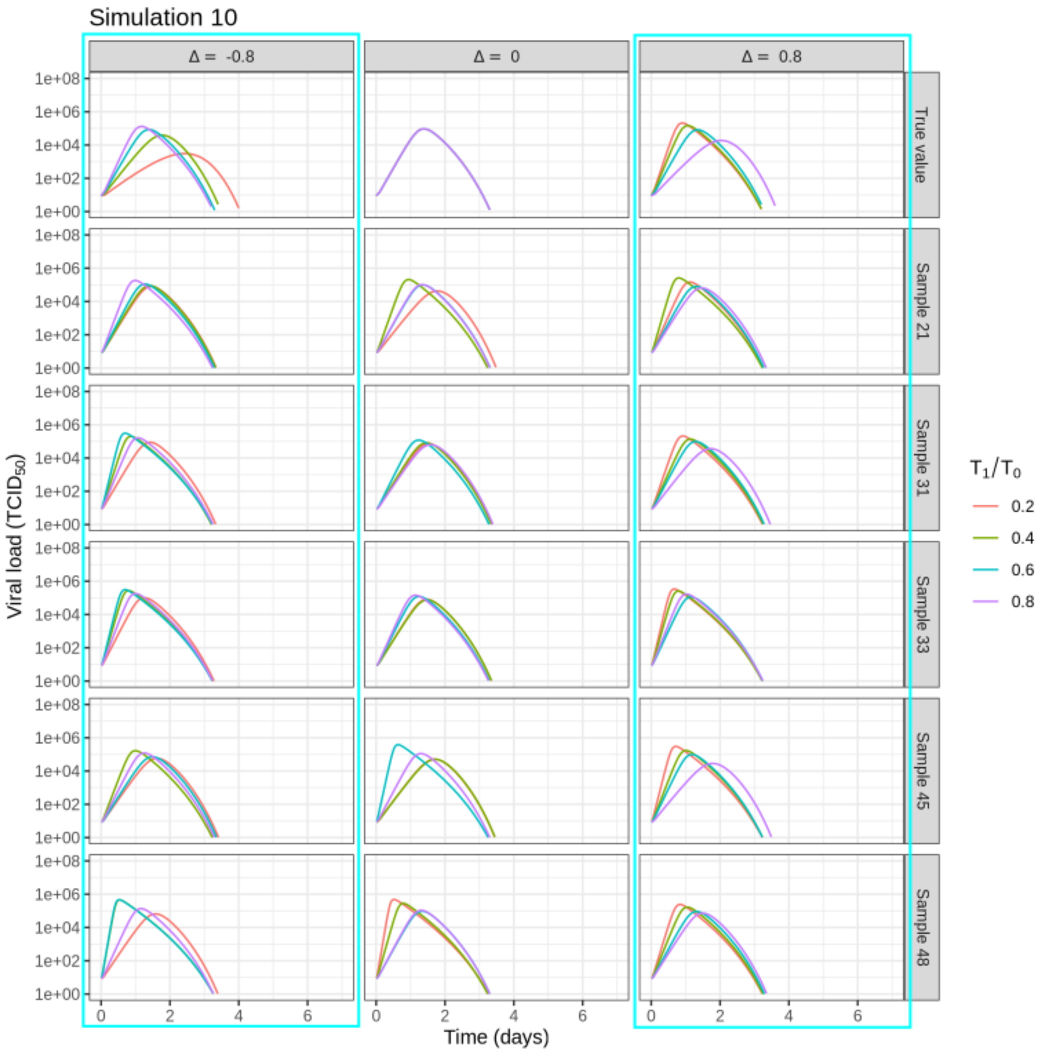


Figure 10: Model simulations with true and inferred parameter values for parameter set 10.
